# Integration of transcriptome, proteome and phosphoproteome data elucidates the genetic control of molecular networks

**DOI:** 10.1101/703140

**Authors:** Jan Großbach, Ludovic Gillet, Mathieu Clément-Ziza, Corinna L. Schmalohr, Olga T. Schubert, Christopher A. Barnes, Isabell Bludau, Ruedi Aebersold, Andreas Beyer

## Abstract

Genomic variation affects cellular networks by altering diverse molecular layers such as RNA levels, protein abundance, and post-translational protein modifications. However, it remains unclear how these different layers are affected by genetic polymorphisms and give rise to complex physiological phenotypes. To address these questions, we generated high-quality transcriptome, proteome, and phosphoproteome data for a panel of 112 genetically diverse yeast strains. While genetic effects on transcript abundances were generally transmitted to the protein level, we found a significant uncoupling of the transcript-protein relationship for certain protein classes, such as subunits of protein complexes. The additional phosphoproteomics data suggests that the same genetic locus often affects distinct sets of genes within each of these layers. In particular, QTLs tended to affect upstream regulatory proteins at the phosphorylation layer, whereas downstream pathway targets were typically affected at the transcript and protein abundance layers. Underscoring the importance of regulatory protein phosphorylation in linking genetic to phenotypic variation is the finding that the number of protein phosphosites associated with a given genetic locus was more predictive for its influence on cellular growth traits than the number of transcripts or proteins.

This study shows how multi-layered molecular networks mediate the effects of genomic variants to more complex physiological traits and highlights the important role of protein phosphorylation in mediating these effects.

## Introduction

Genetic polymorphisms are important modifiers of many physiological traits, such as body height or disease susceptibility. Differences in these traits are caused by alterations of the underlying molecular networks^1^. There are various ways how genetic variants can influence these molecular networks. For instance, genetic variants might either act indirectly through multiple layers, e.g. by first changing mRNA levels, which then influence protein levels, or directly on individual layers, e.g. by changing protein levels through altered protein stability^2^. Changes in the sequence of an mRNA can affect translation rates, hence altering protein abundance, while the concentration of its corresponding mRNA stays constant (reviewed in ^3^). The net activity of proteins can be modulated by changing the protein levels and/or by changing their specific activities, e.g. by increasing or decreasing regulatory phosphorylation. Varying concentrations of a kinase can affect phosphorylation levels of its targets, which, in turn, might lead to their degradation^4^, activation or inhibition^5^, or changes in their subcellular localization^6^. The local concentration of a specific phosphorylated protein (e.g., a transcription factor) might in turn affect transcription of yet other genes. Thus, the crosstalk between the different molecular layers is characterized by a great diversity of mechanisms, including biosynthetic- and signalling processes^2^.

High-throughput molecular profiling (‘omics’) technologies have enabled the investigation of the impact of genetic variation on individual molecular layers on a genomic scale. Such data can be used to identify genetic variants that explain some of the variation in the measured traits, termed quantitative trait loci (QTLs). For example, with current RNA sequencing (RNA-seq) technologies it is possible to detect expression QTLs (eQTLs) for virtually all transcripts present in a cell^7^. Likewise, mass spectrometry has been used to identify QTLs for hundreds of proteins (pQTLs)^8–11^ and metabolic traits (mQTLs; reviewed in ^12^). A shortcoming of many existing studies is that either only one molecular layer was measured, typically transcripts, or, if multiple layers were measured, transcripts and proteins were not isolated from the same cultures, reducing comparability due to potential differences in environmental effects. Furthermore, post-translational protein modifications (PTMs), such as phosphorylation, have been largely neglected and no systematic phosphorylation QTL (phQTL) studies have been performed to date.

A comprehensive view of the effects of genetic perturbations on integrated molecular networks requires accurate quantitative measurements at different molecular layers from the same samples^14^. Therefore, we designed a multi-omics QTL study in recombinant offspring of a cross of two budding yeast strains, with four key distinguishing features: (i) we quantified transcripts, proteins, and protein phosphorylation levels at a genomic scale; (ii) by using the SWATH-MS technology we could reproducibly quantify a large number of proteins across virtually all samples; (iii) RNA, protein, and phosphoprotein samples were obtained from the same yeast cultures, which greatly facilitated the integration of the data; and (iv) the data integration scheme that we developed for this study enabled us to investigate interdependencies of the molecular patterns between the layers. This setup enabled us, for the first time, to map the response of cellular signalling networks to genomic variation, to dissect the interaction of transcript- and protein abundance changes, and to investigate the relevance of phQTLs for complex cellular traits.

## Results

### Multi-omics profiling of a yeast panel

The BYxRM *Saccharomyces cerevisiae* yeast segregant panel used in this study results from a cross of a laboratory strain (BY4716) isogenic to the reference strain S288C, and a derivative of a wild isolate from a vineyard (RM11-1a). This cross was previously used to study the genetic contribution to molecular traits – including RNA and protein levels – and to test novel systems genetics approaches^15–17^. We grew the two parental haploid yeast strains and 110 of their recombinant offspring under tightly controlled conditions, with multiple replicates for some of the strains (Fig. 1, Supplementary Data S1). The transcriptomes of 150 cultures were sequenced at high coverage (38-186x) allowing for the quantification of 5,429 transcripts in all samples (Supplementary Data S2). The RNA-seq data was also used to infer the genotypes of the strains, using a strategy that we developed previously^18^ (Supplementary Figure S1, Supplementary Data S3), significantly improving the localization of recombination sites compared to previous studies^19^. In samples obtained from the same yeast cultures, we used SWATH-mass spectrometry (MS) to measure the abundances of 1,862 proteins with less than 1.8% missing values across all samples (Fig. 2a, Supplementary Data S4). This represents a four-fold increase in the number of quantified proteins compared to previous studies in the same cross^8,11,16,17^. We also quantified the phosphorylation state of the proteins by SWATH-MS; after stringent filtering, we obtained abundances of 2,116 phosphopeptides from 988 proteins with less than 2.4% missing values across all samples (Supplementary Data S5).

**Figure 1:**
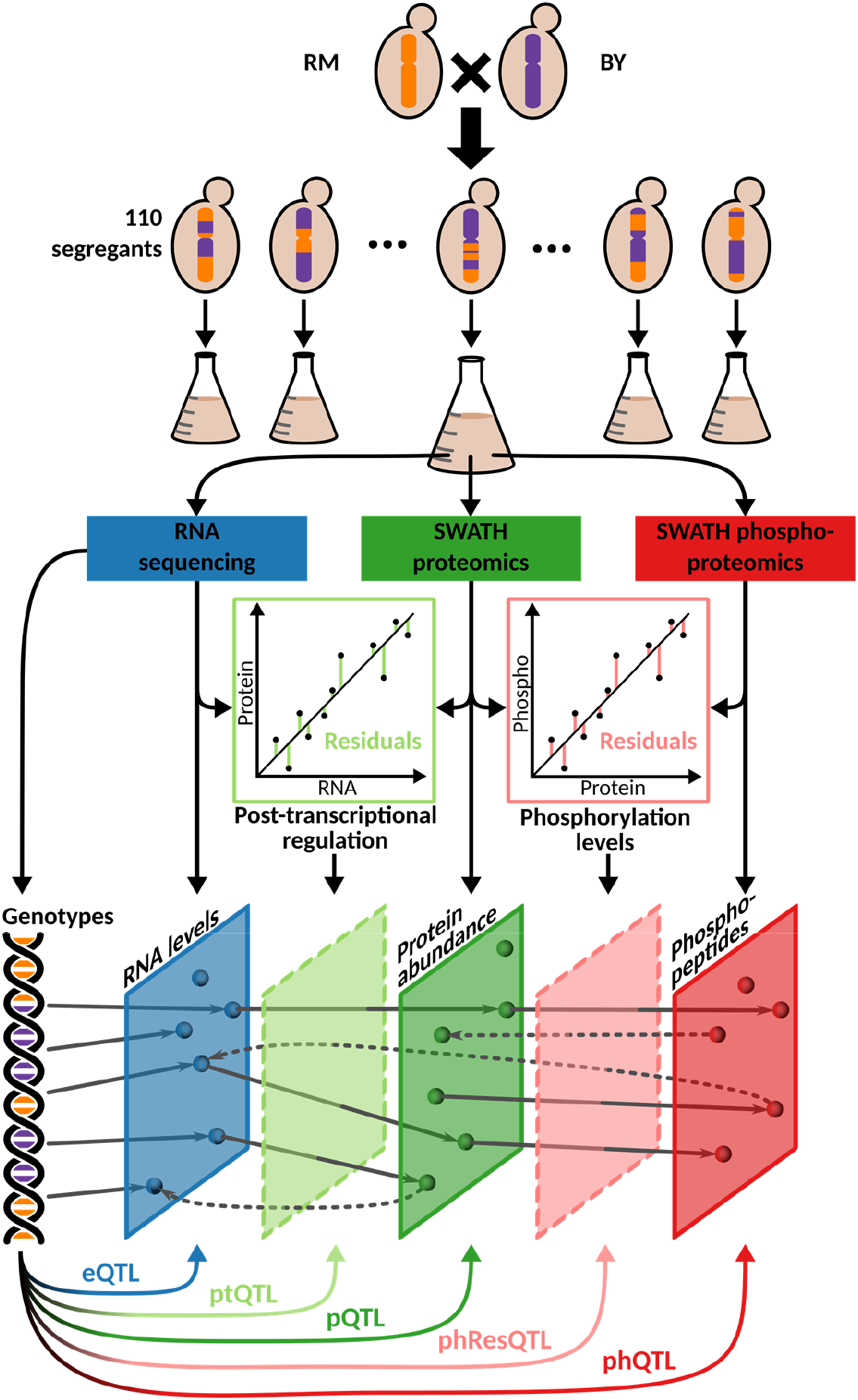
Experimental design and study overview. Yeast segregants derived from two parental strains, BY and RM, were grown and characterized with different omics approaches. Their transcriptome, proteome, and phosphoproteome were directly quantified in samples obtained from the same culture flask. Residual traits representing the disparities between transcript and protein levels (i.e., measuring post-transcriptional regulation, light green), and those between protein and phosphopeptide levels (i.e., measuring phosphorylation status, pink) were computed. QTL analyses were performed to determine the effect of genetic variation on each of these molecular layers.

**Figure 2:**
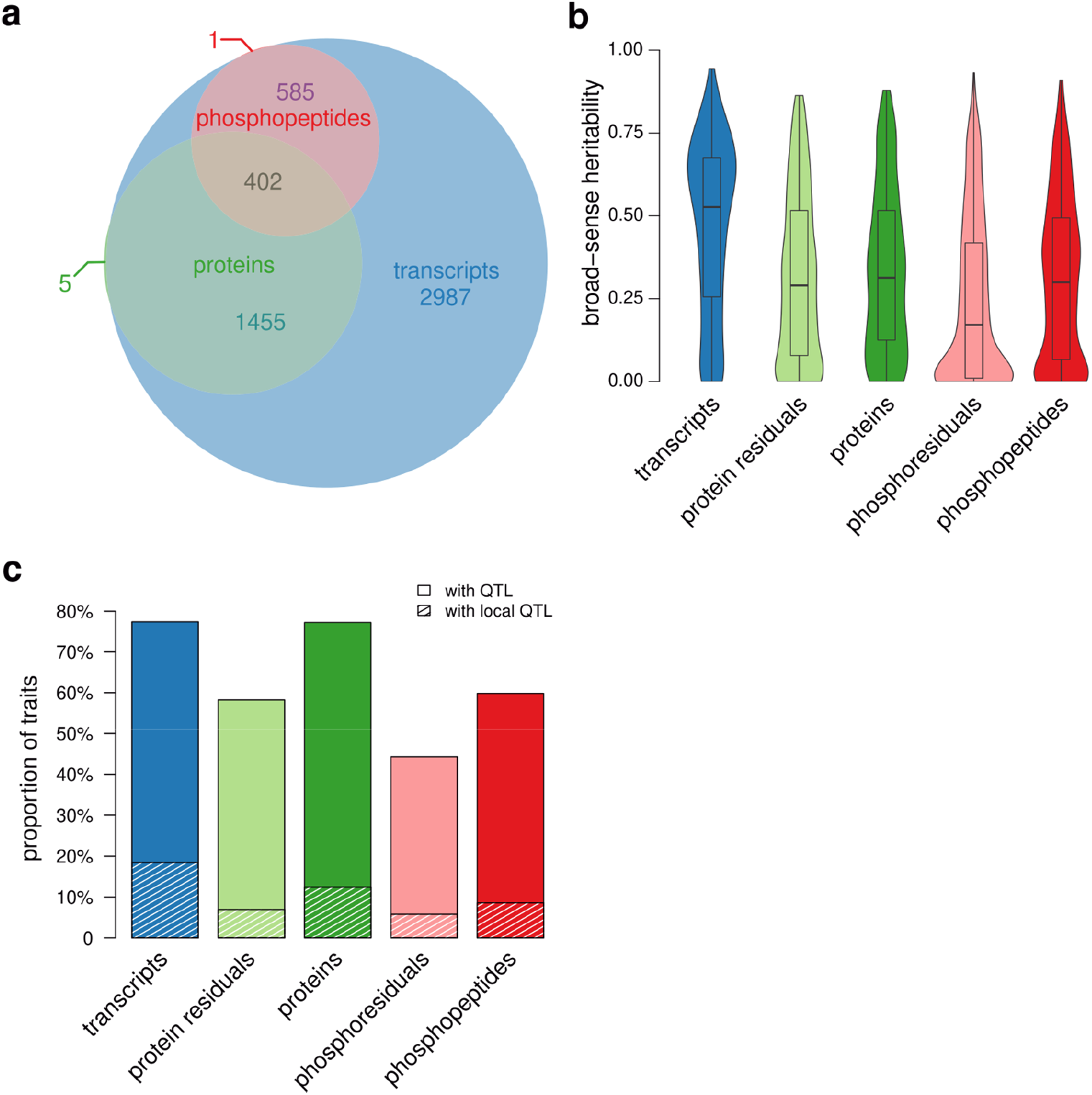
Genetic control of molecular phenotypes. **a**, Overlap of quantified transcriptome, proteome, and phosphoproteome. **b**, Broad-sense heritability of traits belonging to the 402 genes for which measurements at each molecular level were available. Broad-sense heritability was estimated by comparing the variation between replicates from the same strain (which is non-genetic) to the total variation between strains. When the intra-strain variation is small compared to the inter-strain variation, it can be concluded that the genetic contribution to trait variation is large under the experimental conditions tested^20^. **c**, Proportion of molecular traits affected by at least one QTL are shown separately for each molecular layer. Shaded areas indicate the proportion of traits for which a local QTL was found.

As expected, we observed that in most cases, protein abundances were positively correlated with their transcript levels (average r=0.23), and most phosphopeptides were positively correlated with their proteins of origin (average r=0.29) (Supplementary Figure S2). However, each layer can also be affected independently of the genetic effects on the other layers. In order to identify direct effects that act on a molecular layer, we computationally generated traits from which effects acting on other molecular layers were removed as previously proposed^8^. First, we estimated the contribution of mRNA changes to protein-level changes by regressing the concentration of a given protein against the concentration of its encoding mRNA across all strains. Deviations from this regression (residuals) can either result from noise in the data, or from pQTL effects that are independent of transcript-level changes. We used the residuals as estimates of post-transcriptional regulation, resulting in 1857 post-transcriptional (pt) traits^8^. Likewise, we regressed phosphopeptide levels against levels of the proteins of origin and used the residuals as estimates of differential phosphorylation, resulting in 879 phospho-residual (phRes) traits. The QTLs obtained by mapping these residual traits were termed post-transcriptional QTL (ptQTL) and phospho-residual QTL (phResQTL), respectively.

### QTLs have widespread effects on all quantified molecular layers

In order to quantify the fraction of trait variation that can be attributed to genetic differences, we quantified broad-sense heritability by leveraging available replicates as proposed before^20^ (Fig. 2b). All five types of molecular traits outlined above had heritabilities greater than expected by chance (p<2.2E-16, Wilcoxon rank sum test); 525 (28%) pt traits and 165 (18%) phRes traits had heritabilities greater than 50%, indicating that at least a part of the resi dual variation isgenetically determined and not just technical and/or biological noise^8^.

We utilized a Random Forest-based mapping strategy to identify QTLs. This approach was previously shown to outperform traditional QTL mapping methods because of its capability to account for complex (epistatic) interactions between genetic loci^21^. In total, we detected 5,776 eQTLs, 2,078 pQTLs, and 1,327 ptQTLs at a false discovery rate (FDR) below 10% (Supplementary Table S1, Supplementary Figures S3a-e). The same fraction of transcripts and proteins had at least one QTL (77% at FDR<10% in both cases; Fig. 2c). This large proportion of proteins with at least one pQTL underlines the high quality of the proteomics data^8^. We also detected 1,595 phQTLs and 466 phResQTLs affecting 1,266 phosphopeptides (60%) and 389 phosphoresiduals (44%), respectively (Supplementary Table S1).

To understand where these QTLs are located with respect to their target genes, we classified QTLs as either local or distant based on their linkage disequilibrium with the genetic marker that is closest to the affected gene (Methods). The fraction of molecular traits with a local QTL was in a similar range for all directly measured traits (10-20%), with transcripts being most strongly enriched for local QTLs (Fig. 2c). The residual-derived traits (pt and phRes) had the smallest fraction of local QTLs, which may be due to biological reasons, due to increased noise (as discussed above), or a combination of the two. Next, we asked whether local eQTLs and pQTLs could be attributed to changes in the sequence of the respective transcript, which might influence transcription and translation rates. When comparing genes with local QTLs (i.e., local eQTLs, pQTLs, and ptQTLs) with genes that are only affected by distant QTLs, we found that the former had an increased number of polymorphisms in non-coding regions (e.g., 5’ untranslated regions (UTRs) or 3’ UTRs; Fig. 3a). The existence of local ptQTLs (129 local ptQTLs for 6.9% of all pt traits) and the fact that ptQTLs were enriched for polymorphisms outside of coding regions, suggest that variants in non-coding parts of the genome can influence protein levels independently of their coding transcripts^8^, for example through polymorphisms in ribosomal binding sites affecting translation initiation or variants altering mRNA capping and looping.

**Figure 3:**
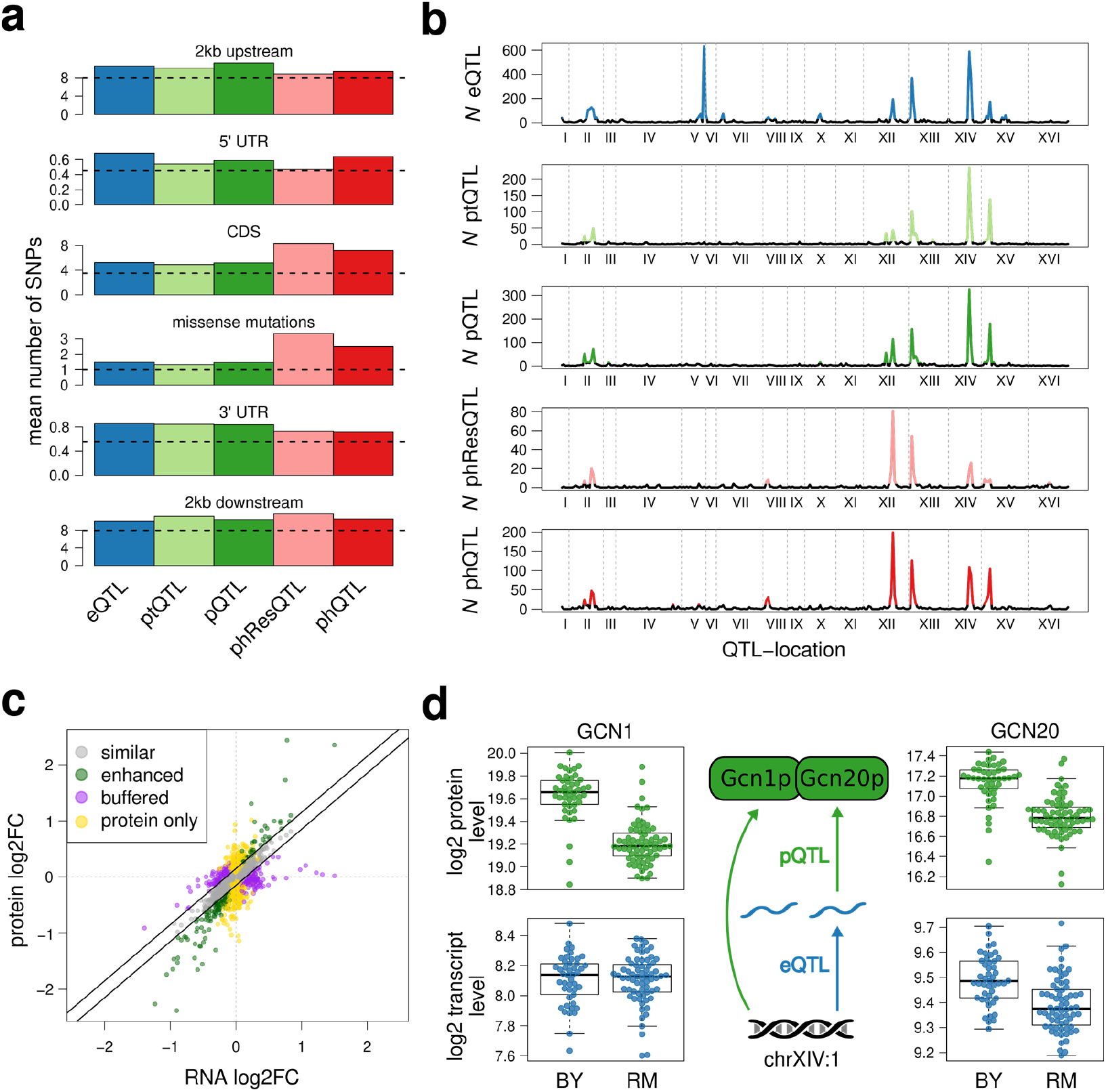
Genomic effects on transcript and protein levels. **a**, Average number of SNPs in and around genes affected by local QTLs at the respective molecular layer. The horizontal dashed black line shows the average number of SNPs across all annotated genes in the respective regions (Methods). **b**, Number of traits affected by each locus at FDR<10% for each molecular layer. Regions with more QTLs than expected by chance (QTL hotspots) are shown in color. **c**, Effects of QTLs on transcript and protein levels. Each dot represents an association of a QTL with a gene. The relationship between effect size and direction on the transcript and protein levels are color-coded based on the four effect classes described in the main text. Axes show the log2-transformed fold changes of BY versus RM alleles. **d**, Effects of the hotspot chrXIV:1 on the Gcn1-Gcn20 protein complex (y-axis shows log2-transformed read counts and protein abundances). Trait levels for each strain are shown separately for each allele as dots for each gene/protein. *GCN20* is affected at the transcript and protein levels while the transcript levels of *GCN1* did not change significantly.

Regions in the genome that affect significantly more traits than expected by chance are referred to as QTL hotspots^22^. Since these loci affect large numbers of traits, they are assumed to act through master regulators such as transcription factors or kinases^7,23^. We tested for regulatory hotspots as proposed before^15^ (Methods) and detected between 9 and 15 significant hotspots for each molecular layer, with the largest number of hotspots being detected for the eQTL layer (Fig. 3b, Supplementary Table S2). Previous work found that most distant eQTLs act from within hotspots^7^. Our data shows that this is the case for QTLs of all five types of molecular traits considered here (Supplementary Figure S4). Many of the detected hotspots have been reported as eQTL hotspots for this yeast cross before. For some of them a causal gene has been validated, e.g., *HAP1* (chrXII:2), *IRA2* (chrXV:1), and *MKT1* (chrXIV:1)^15,22^. While most of the hotspots affected multiple molecular layers simultaneously, we also observed hotspots that predominantly impacted the transcriptome (e.g., chrV:2), the proteome (e.g., chrXII:1) or the phosphoproteome (e.g., chrVIII:1). Thus, the molecular mechanism of a QTL may have a strong influence on which molecular layer is primarily affected.

### Transmission of genetic effects from the transcriptome to the proteome

The deep coverage of the transcriptome and proteome in this study enabled us to investigate to what extent transcript level changes are transmitted to their corresponding proteins. First, we observed that local eQTLs and local pQTLs were significantly overlapping (p<2.2E-16, OR=11, one-sided Fisher’s exact test), which is expected in the absence of major post-transcriptional or post-translational regulation. In order to estimate QTL effect sizes, we split the population of yeast segregants based on the alleles at a linked locus and computed the log fold change of the transcript and protein levels between the two sub-populations. When comparing transcript fold changes with the respective protein fold changes, we found that they were strongly correlated (eQTLs with FDR<10%, r=0.73, Fig. 3c). Previous work suggested that local and distant eQTLs affect the proteome in different ways (^8,13^, Supplementary Text S1). Using our data, a regression of transcript and protein fold changes had a slope close to 1 for both local and distant eQTLs (Supplementary Figure S5), implying that changes in transcript levels tend to cause similar changes in protein levels regardless of the eQTL being local or distant. In addition, for most eQTL hotspots, transcript variation was propagated to protein levels, as for example at the *HAP1* locus (chrXII:2). Effects of this hotspot on protein concentrations were highly correlated with those at the transcript level (Pearson’s correlation coefficient r=0.94 for the 289 eQTLs at FDR<10%).

Despite widespread concordance between eQTL effects on transcripts and proteins, we also detected many eQTLs exhibiting effects on the protein level that were different from those on the transcript level. We classified targets of eQTLs into three groups based on the difference of allelic effects at the transcript and protein levels (Methods and Fig. 3c). The first group contained genes for which the effects of an eQTL on the transcript and protein levels of its target gene were similar (‘similar’). The second group contained genes for which the effects of an eQTL were repressed or even entirely buffered on the protein level (‘buffered’). The third group contained genes for which proteins showed enhanced responses compared to their corresponding transcript (‘enhanced’). As a fourth group, we added genes that were affected on their protein level by ptQTLs, without a significant effect on the corresponding transcript level (‘protein only’). Gene Ontology (GO) enrichment revealed that genes with eQTLs that affected their protein level to a similar extent as the transcript levels were enriched for genes related to protein import into the mitochondrial matrix (Supplementary Table S3). Genes with buffered eQTL effects were strongly enriched for terms related to cytoplasmic translation, including the large subunit of the ribosome, and proteins localizing to the nucleolus (Supplementary Table S4). Although buffering of ribosomal proteins has been observed before^8^, it could not be excluded that this was due to technical issues in protein quantification. However, this is unlikely the case here: first, because ribosomal proteins are relatively highly expressed and hence easily quantifiable by mass spectrometry, and second, because these proteins were affected by other pQTLs at a similar rate as the rest of the proteome (73% of proteins with buffered eQTLs had at least one pQTL).

Genes affected by enhanced eQTLs were strongly enriched for mitochondrial ribosomes and other terms related to mitochondrial translation (Supplementary Table S5). Like buffered proteins, proteins affected by ‘protein only’ effects were also enriched for functions related to cytoplasmic translation (Supplementary Table S6). This unexpected functional similarity between buffered proteins and proteins subject to ‘protein only’ effects raised the question whether the same proteins could be subject to both phenomena. Indeed, we found that genes affected by a buffered eQTL were more likely to also be affected by a ‘protein only’ QTL (219 genes; p<7E-4, Fisher’s exact test). Thus, specific groups of proteins seem to require extensive post-transcriptional fine-tuning of their cellular concentrations, decoupling protein levels from transcript levels (Supplementary Figure S6).

This notion was further supported by detailed investigation of individual hotspots with heterogeneous eQTL and pQTL effects. For example, the effects of the *IRA2* hotspot (XV:1) on transcript levels of cytoplasmic ribosomal genes was not transmitted to protein levels, but the same locus affected protein levels of genes related to mitochondrial respiration without changing their transcript levels (Supplementary Figure S7). While the effects of the *MKT1* hotspot (XIV:1) on the protein levels of cytosolic ribosomes were buffered, the effects of the same locus on the protein levels of mitochondrial ribosomes were enhanced. Thus, our analysis reveals complex post-transcriptional QTL effects, especially for genes involved in translation, with marked differences between cytoplasmic and mitochondrial translation.

### Protein complex stoichiometry controls QTL effects

The buffering of cytoplasmic ribosomal proteins is a well-described phenomenon: excess proteins not incorporated into ribosomes get degraded^24^. Recently, buffering of components of other protein complexes was reported^25–27^, suggesting that the need to maintain protein complex stoichiometry drives post-transcriptional protein abundance regulation. Indeed, we observed extensive co-regulation of genes whose products participate in common complexes on multiple molecular layers: both transcript levels and protein levels of protein complex members were correlated across the strains (average Pearson’s correlation on RNA and protein levels: r=0.8 and r=0.46, respectively). Furthermore, pairs of genes of the same complex were affected by the same eQTL or pQTL at a significantly higher rate than pairs of genes from different complexes. Whereas this phenomenon was particularly strong for ribosomal proteins, we still observed significant enrichment after excluding ribosomal proteins from the analysis (eQTLs: p<7E-13, OR=3.45, Fisher’s exact test; pQTLs: p<7E-9, OR=3.06). Moreover, proteins in the same complex were even more frequently affected by the same ptQTL (p<2.2E-16, OR=6.18), indicating that their levels are co-regulated independently of their RNA levels. Indeed, we observed that most proteins from complexes correlated better with other proteins in the same complex than with their own transcript (75%, Supplementary Figure S8), confirming that the stoichiometry of complexes is a major driver controlling protein levels. One example for this is the Gcn1p-Gcn20p complex, which plays a role in translation initiation. Whereas both proteins were affected by a common pQTL, only *GCN20* transcripts, but not *GCN1* transcripts were affected by the respective eQTL (Fig. 3d, ^28^). Hence, the protein level effect on Gcn1p is likely an indirect response to the Gcn20p change, thereby re-establishing stoichiometric ratios.

Taken together, our data supports protein complex stoichiometry as an important post-transcriptional effector of protein abundance that substantially contributes to ptQTL effects.

### Protein phosphorylation is often regulated separately from protein levels

To investigate effects acting directly on the phosphorylation state of proteins, we focused on the phRes traits for which the phosphorylation effects were corrected for abundance changes of the proteins of origin. After this correction we still detected 466 phResQTLs (44% of all phRes traits), including multiple phResQTL hotspots (Fig. 3b). A striking example of a phResQTL hotspot is the *HAP1* locus, affecting 22.1% of all phosphoproteins, but only 6.5% and 5.2% of the corresponding transcripts and proteins of the 402 genes whose products were detected on all three molecular layers (Fig. 4a).

**Figure 4:**
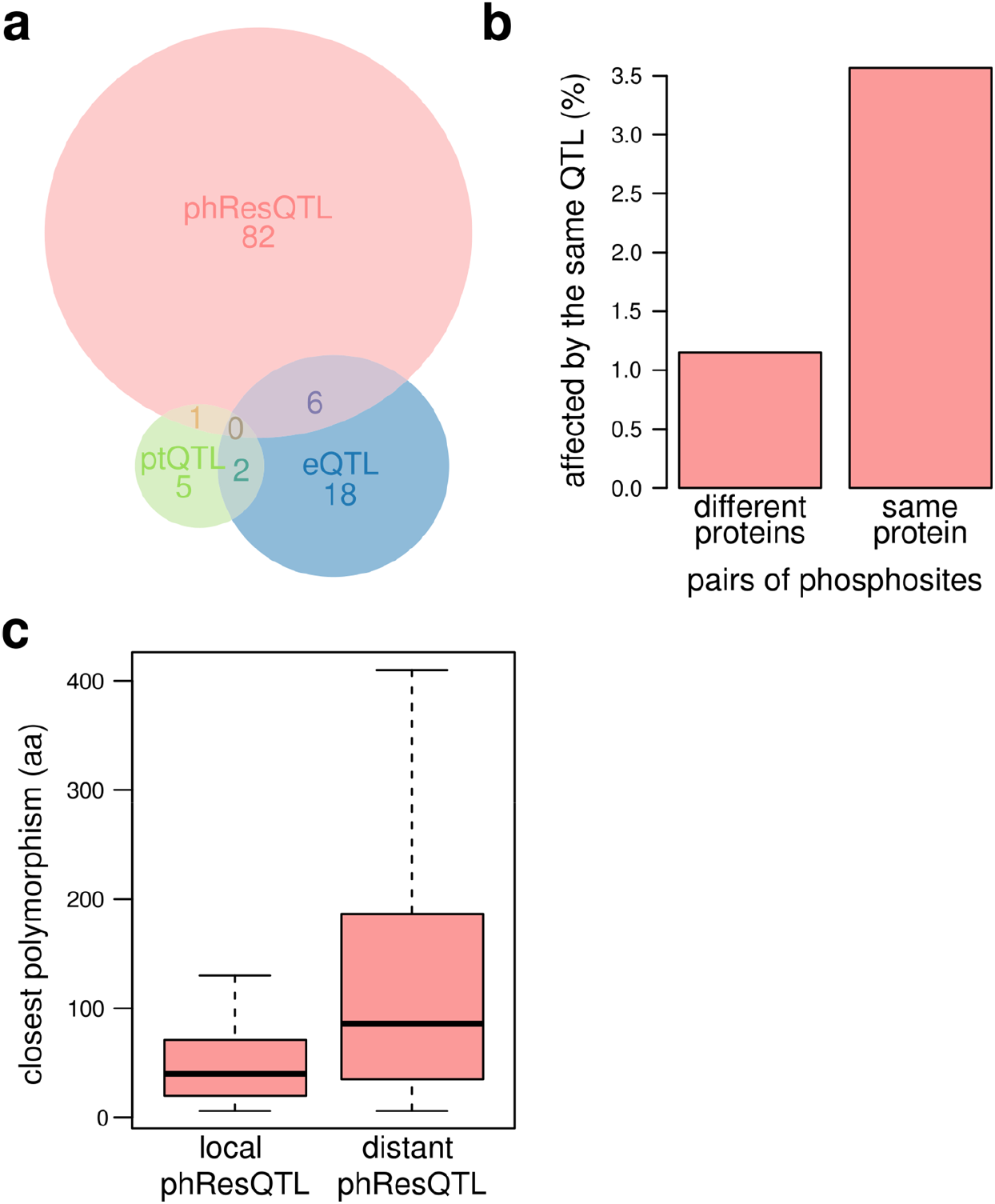
Analysis of phResQTLs. **a**, Overlap of different QTL types in the *HAP1* locus (chrXII:2) for the 402 genes whose products were detected on the transcript, protein, and phosphopeptide level. **b**, Proportion of pairs of phosphosites on different or the same protein that share a phResQTL. **c**, Distance of phosphosites affected by a local phResQTL or exclusively by distant phResQTLs to the nearest missense mutation in the protein of origin.

Next, we asked whether genomic variation could result in coordinated effects on different phosphosites on the same protein. Indeed, phRes traits of the same protein (i.e., different phosphosites on the same protein) had a higher chance to be targeted by the same phResQTL than random pairs of phosphosites (p<3E-8, one-sided Fisher’s exact test; Fig. 4b). This is consistent with the notion that multiple phosphosites on the same protein are often targeted by a common kinase or phosphatase^29^.

Local phResQTLs might be caused by genetic variants directly affecting the phosphorylation state of a protein, for example through the modification of residues close to a kinase or phosphatase binding site. In support of that notion, we found that proteins that were affected by at least one local phResQTL had more missense polymorphisms than proteins that only had distant phResQTLs (p<1.3E-07, one-sided Wilcoxon rank sum test; Supplementary Figure S9). In addition, phosphosites with a local phResQTL were closer to a missense variant than those with a distant phResQTL (p<2.4E-05, Fig. 4c). Further, if two phosphosites on the same protein were both affected by two different phResQTLs – one local and the other distant – the phosphosite with the local phResQTL was on average closer to a polymorphism than the distant one (p=0.005, one-sided paired Wilcoxon rank sum test; Supplementary Figure S10). In summary, these results suggest that many local phResQTLs are caused by missense polymorphisms, changing recognition motifs or 3D structures of affected proteins, and thereby affecting binding of kinases and phosphatases to such phosphosites.

### Phospho-QTLs are strongly associated with cellular fitness effects

In order to better understand the mechanisms by which genetic variation affects cellular fitness we integrated our molecular QTL data with growth traits in 46 different conditions measured earlier for the same yeast cross, including differing temperatures and pHs, multiple carbon sources, and exposure to metal ions and small molecules^20^. We found that QTL hotspots affecting molecular traits were more likely than other genomic regions to also affect growth rates (Supplementary Figure S11a,b, Supplementary Table S7), which underlines the contribution of hotspots to complex growth trait variation. Importantly, the number of growth traits linked to a hotspot was best predicted by the number of phosphorylation traits targeted by the same hotspot, whereas all other molecular layers were less predictive (Supplementary Figure S11c). This unique role of a post-translational response to genetic variability warranted a closer inspection of state changes in regulatory networks. To study QTL effects on the ‘regulome’, we integrated the five types of molecular traits with regulatory network information. We found that phResQTL targets were often functionally related to the putative causal genes underlying a given hotspot and/or targets of kinases that were affected by the same locus. The *IRA2* hotspot^22^ is an example for such a case. This hotspot affected 36 growth traits and had targets among all five types molecular traits (Supplementary Table S2, Fig. 3b). Ira2 inhibits the Ras/Pka pathway by promoting the GDP-bound form of Ras2, which is crucial for the adaptation of cellular metabolism to conditions with different nutrient availabilities ^30^. Earlier work has shown that polymorphisms in the *IRA2* coding sequence affect the activity of the Ras/Pka pathway and that the RM allele of *IRA2* inhibits this pathway more efficiently^22^. Several targets of phQTLs in the *IRA2* hotspot were functionally connected to *IRA2* including a local phQTL targeting Ira2 itself, a phQTL targeting Cdc25, which promotes the GTP-bound form of Ras2, and a phQTL targeting the Ras2 protein^31^. In addition, we detected phQTLs for numerous downstream targets of Ras/Pka signaling at the *IRA2* hotspot (Supplementary Table S8). Hence, our data reveals widespread modifications of key signaling molecules in the Ras/Pka pathway upon genetic variation and contributes to a better understanding of the molecular mechanisms through which this QTL hotspot acts.

Another example illustrating the effect of genetic variants on cellular signaling is the *HAP1* hotspot on Chromosome 12 (chrXII:2)^15,17^, which was previously shown to affect 19 growth traits^20^. Hap1 has been shown to be involved in the regulation of respiration in response to oxygen and iron deprivation^32^. Again, our analysis revealed that this locus has much more prominent effects on the phospho-layer than on transcript or protein abundances (Figure 4a). The integration of our multi-layered QTL data with previously published kinase-substrate networks suggested a new mode of effect for this hotspot: both the expression and phosphorylation of Psk2, a protein kinase and known regulator of carbohydrate metabolism^33^, were significantly regulated by the *HAP1* hotspot. Proteins whose phosphorylation was affected by the *HAP1* locus were enriched in previously reported targets of the kinase Psk2^34^. This enrichment of phosphorylated Psk2 substrates strongly implies a directed signaling cascade, from the *HAP1* locus *via* altered Psk2 activity to the Psk2 substrates.

The added value of integrating protein phosphorylation with transcript and protein abundances was even more apparent from a hotspot on Chromosome 8 (chrVIII:1) with the pheromone response gene *GPA1* harboring the causal mutation^23^. The hotspot harbored only 69 eQTLs, 8 pQTLs, and 1 ptQTL, but 41 phQTLs (Supplementary Table S2, Fig. 3b). Many of the hotspot eQTL targets (41%) could be found downstream of the mating pheromone response pathway, which supports *GPA1* as a causal gene. However, several targets of this hotspot, especially those on the phospho level such as Rck2 and Rsc9, were involved in the osmotic stress response, which is unrelated to *GPA1* ^35–37^. This suggests that the polymorphisms in *GPA1* do not fully explain the effects of this hotspot, prompting us to look for other potential key effectors of the hotspot. The genetic marker with the most phQTL targets within the hotspot is located within the coding region of *STE20*. Ste20 is the key activator of multiple mitogen-activated protein kinase (MAPK) pathways, including the mating pheromone response, but also invasive growth regulation, regulation of sterol uptake, and, importantly, osmotic stress response ^35^(Fig. 5). Indeed, for up to 60% of the target genes of this hotspot we found a link to Ste20 and/or Gpa1 (Supplementary Tables S9 and S10). Further, most of the phosphopeptides and phRes traits targeted by the hotspot corresponded to proteins phosphorylated by components of MAPK pathways downstream of Ste20 (51%; Fig. 5, Supplementary Tables S9 and S10). GO enrichment analysis of targets of the hotspot at the transcript layer revealed an enrichment for biological processes under the influence of *STE20* (Supplementary Table S11). Taken together, these results suggest that the effects of the chrVIII:1 hotspot are due to the combined effects of the polymorphisms in *GPA1* and *STE20*. Furthermore, we provide new potential components of the MAPK pathways that are regulated by this hotspot but were not previously identified as downstream targets of Ste20 (Supplementary Table S12).

**Figure 5.**
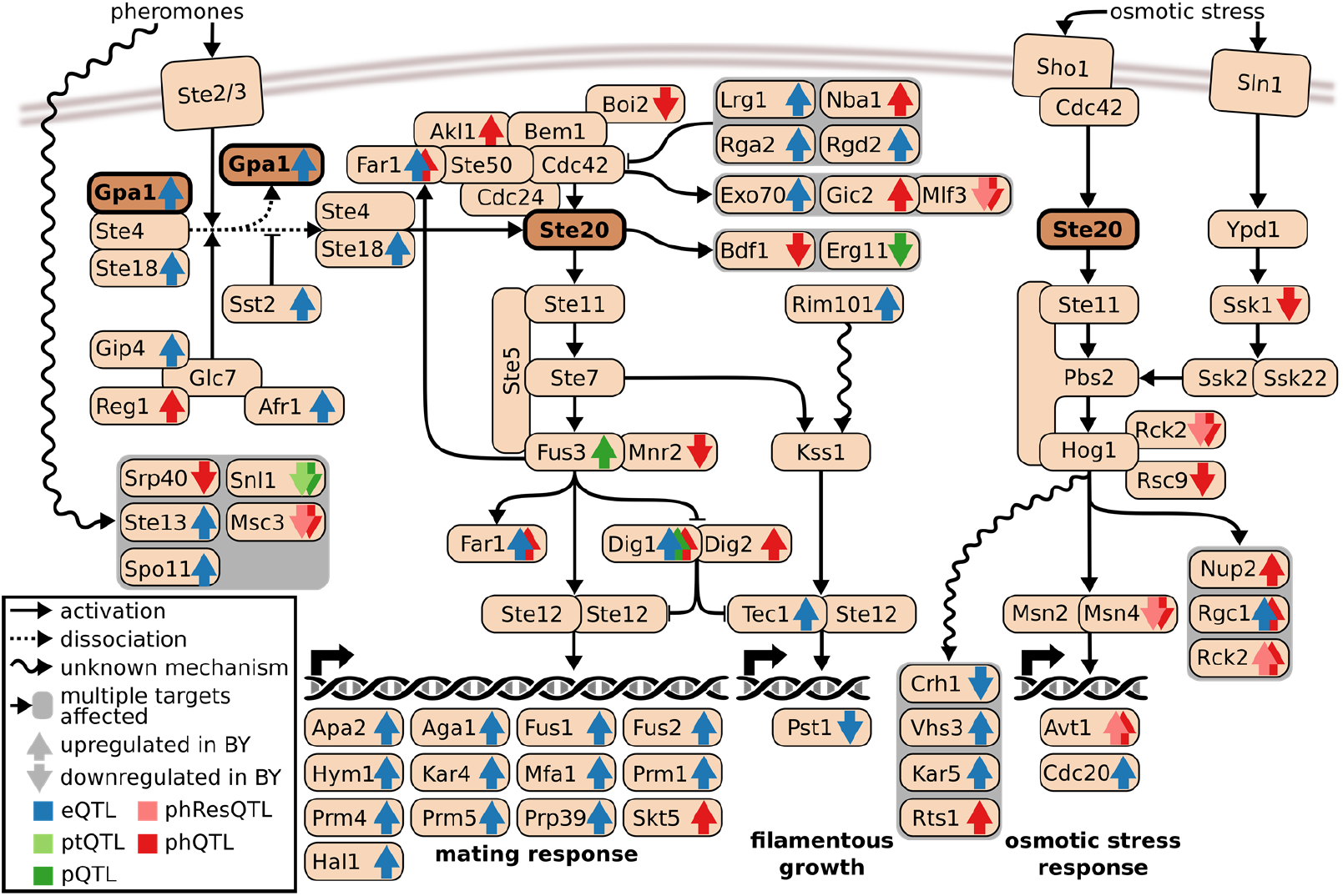
Multi-layer effects of the hotspot around *GPA1* and *STE20* on Chromosome 8. Gpa1 and Ste20 play key roles in the mating response pathway (left, through transcription factor complex Ste12/Ste12) and filamentous growth regulation pathway (middle, through transcription factor complex Tec1/Ste12). Ste20 additionally regulates the osmotic stress response pathway (right, through transcription factors Msn2/Msn4). Depicted are genes that are known to have a direct or indirect connection with *GPA1* and/or *STE20*. Genes that we found to be affected by the *GPA1*/*STE20* hotspot on any molecular layer are indicated with arrows in the respective color of each layer. The arrow orientation indicates the fold change direction. References for each known connection are given in Supplementary Table S9.

The examples of these hotspots illustrate how phQTL mapping provides information on signaling networks that is orthogonal to transcript and protein abundance data: genetic variants often affect the phosphorylation states of gene products that are distinct from the genes affected by abundance changes (i.e. eQTL and pQTL targets). Furthermore, those protein activity changes often act upstream of QTL abundance effects. Overall, we show that by quantifying the effects of sequence polymorphisms on multi-layer molecular networks our integrated approach can provide clues toward reconstructing the molecular architecture underlying complex traits and the chain of causality through multiple layers of regulation.

## Discussion

In this study, we present the first dataset that concurrently quantifies the effects of natural genomic variation on the transcript (eQTL), protein (pQTL), and protein phosphorylation (phQTL) layers. Whereas previous ‘omics’ QTL studies have primarily focused on the *abundance* of biomolecules, our study integrates abundance with the *states* of biomolecules. Hence, the conceptual advance of our study is the possibility to discover the consequences of genomic variation, for example the phosphorylation-mediated activation of a signaling pathway component, through the different layers of a molecular network. The three hotspots containing *IRA2*, *HAP1*, and *GPA1*/*STE20* exemplify this notion.

Quantifying transcript and protein levels along with the state of the proteins indicated by phosphorylation from the same yeast culture maximized the comparability of the data, thus facilitating data integration. This three-layer molecular profiling enabled the following findings: (i) RNA effects are generally transmitted to protein levels in a 1:1 relationship. (ii) There are, however, numerous exceptions and specific classes of genes are subject to enhanced (e.g., mitochondrial ribosomes) and buffered (e.g., cytosolic ribosomes and nucleolus) protein-level effects. (iii) The maintenance of protein complex stoichiometry plays a major role in the post-transcriptional effects of genetic variants. (iv) A substantial fraction of phosphopeptides (44%) were under the direct control of at least one QTL, independently of the abundance of its protein of origin. Further, we observed multiple cases in which protein abundance effects were buffered (i.e., repressed) at the level of phosphopeptides. (v) QTLs affect multiple phosphosites on the same protein more often than expected by chance. (vi) Protein phosphorylation is much more often affected by local missense variants than protein levels and, thus, the structural destabilization of proteins through segregating variants seems to be a rare event and is presumably under strong negative selection. (vii) Variants affecting phosphorylation are typically in close proximity to the affected residue. (viii) The characterization of phosphorylation events in addition to transcript and protein abundances enabled the separation of upstream and downstream effects of genetic variants on signaling pathways. Importantly, the same QTL can affect distinct sets of genes at different molecular layers. (ix) QTL hotspots were rarely explained by variants directly affecting the transcript encoded at that location; instead, our data suggests that altered protein phosphorylation contributes significantly to the causal mechanisms of several QTL hotspots by the activation of pleiotropic downstream effects.

This study captures some of the enormous complexity in how genetic variants influence signal transmission between different molecular layers, despite all the data being obtained from steady-state standard growth conditions. Clearly, mapping phosphorylation states will be even more critical when aiming to understand the interaction between genetic variants and environmental cues. In addition, future work should address additional aspects of molecular state changes, including other PTMs, changes in protein folding, protein complex formation, and protein-ligand interactions. Our findings have implications for future work on complex human diseases. First, our study shows that, at least in yeast, quantifying phosphoproteomes at large scale and with high precision and reproducibility is feasible with state-of-the-art proteomics technologies. Thus, genome-wide association studies (GWAS) on phosphoproteome changes in human cells might be possible soon as well. Second, our findings imply that mapping phospho-traits and molecular states in general will greatly contribute to understanding the mechanisms through which GWAS loci act. Third, phosphorylation states are more likely to harbor information about physiological processes than mere abundances of transcripts or proteins (Supplementary Figure S11c). Lastly, our results suggest that causal *cis*-variants are often nearby the altered phosphosite, which can help prioritizing candidate variants.

In conclusion, our study sets the stage for a more mechanistic understanding of genomic effects on multi-layered cellular networks and physiological traits.

## Acknowledgements

JG and CS received funding from the German Science Foundation (DFG CRC 680 & CRC 1310). MCZ and CS received funding from the German Federal Ministry for Science and Education (BMBF) as part of the e:Med program (grant no. 01ZX1406). OTS received funding from the German Academic Exchange Service (DAAD), the Swiss National Science Foundation, and the Human Frontiers Science Program. IB was supported by the Swiss National Science Foundation (grant no. 31003A_166435). We thank Dr. Rachel Brem for providing the strains of the BY/RM cross.

## Author Contributions

AB, RA, and MCZ envisioned the study. LG, MCZ, and CAB performed the experiments. JG, CLS, MCZ, OTS, IB, and LG performed the data analysis. All authors contributed to the writing of the manuscript.

## Materials and methods

### Sample preparation

All media were prepared in a single batch to limit experimental variability. The BYxRM yeast strain collection, which we obtained from Rachel Brem, was originally derived from a cross between the two parental strains, BY4716, an S288C derivative (MATα lys2∆0), and RM11-1a (MATa leu2Δ0 ura3Δ0 ho::KAN) ^15^. A subset of 129 strains were picked in random series of 16 (see Supplementary Data S1), pre-cultured in in-house made synthetic dextrose medium (S.D., containing per liter: 1.7 g yeast nitrogen base without amino acids (Chemie Brunschwig), 5 g ammonium sulfate, 2% glucose (w/v), 0.03 g isoleucine, 0.15 g valine, 0.04 g adenine, 0.02 g arginine, 0.02 g histidine, 0.1 g leucine, 0.03 g lysine, 0.02 g methionine, 0.05 g phenylalanine, 0.2 g threonine, 0.04 g tryptophan, 0.03 g tyrosine, 0.02 g uracil, 0.1 g glutamic acid and 0.1 g aspartic acid), and then grown in 115 ml fresh S.D. medium at 30°C until a maximal optical density at 600 nm (OD600) of 0.8 (+/− 0.1). In total, 180 cell cultures were successfully grown to OD600 of 0.8 and then subdivided as follows for the transcript and proteomic analyses respectively. Of the 115 ml of cultures, 15 ml were collected, centrifuged at 2000 g for 3 min at 4°C, transferred into an Eppendorf and snap frozen in liquid nitrogen for transcriptomic analysis. The remaining 100 ml were processed for proteomic analysis essentially as described in ^34^. In short, 6.66 ml of 100% trichloroacetic acid (TCA) was added to the 100 ml culture media to a final concentration of 6.25% and the cells were harvested by centrifugation at 1500 g for 5 min at 4 °C and washed three times with cold acetone. The cell pellets were transferred into 2-ml Eppendorf tubes and frozen in liquid nitrogen.

### RNA sequencing

#### RNA extraction

Total RNA was isolated from deep frozen aliquots of yeast pellets using the RiboPure™ RNA Purification Kit, yeast (Ambion), which includes a DNase treatment to eliminate contamination. RNA quality was assessed using RNA ScreenTape assay (Agilent). All RNAs were of very high quality (Supplementary Data S1, median RIN 9.8, minimal RIN 9.1).

#### RNA-seq library preparation

cDNA libraries were prepared from poly(A) selected RNA applying the Illumina TruSeq protocol for mRNA using a total of 1 μg RNA per sample and 14 PCR cycles.

#### Sequencing

The cDNA libraries were sequenced on a HiSeq2000 with 20 samples per lane (8 million reads per sample). The generated reads were stranded, single-end, and had a length of 100 bp.

### RNA-seq based genotyping

The BYxRM yeast cross has been widely used for QTL mapping. Microarray-based genotype information of the segregants is available^19^. However, deep sequencing enables a more accurate genotyping of recombinant lines. To this end we have exploited published resequencing data of the parental strains^20^ together with the RNA-seq data generated here to infer the genotype of the segregants of the BYxRM cross using a method that we previously developed^18^.

#### Resequencing data of the parental strains

Sequence variation information of the parental strains was obtained from http://genomics-pubs.princeton.edu/YeastCross_BYxRM/; only calls with a MQ≥30 where considered, which represents 42,769 polymorphic sites.

#### Genotype inferring from RNA-seq data

RNA-seq data were mapped to the *S.cerevisiae* reference genome (SaCer3) using TopHat 2 (Ref. ^38^) with the following options: --`min-intron-length 10 --min-segment-intron 10 --b2-very-sensitive --max-multihits 1 --library-type fr-secondstrand`. Read group information was added, and BAM files were sorted using Picard utilities (http://broadinstitute.github.io/picard/).

RNA-seq data were further processed using the GATK pipeline (version 3.4-46-gbc02625) following the best practice guide^39^. First reads containing exon-exon junction were split using the following options: `-T SplitNCigarReads -rf ReassignOneMappingQuality -RMQF 255 -RMQT 60 -U ALLOW_N_CIGAR_READS`, then the variants were called using the UnifiedGenotyper at the 42,769 polymorphic sites identified in the resequencing data of the parental strains. Genotype calls where the GATK genotype score was below 40 (GQ≤40) or that were covered with less than 5 reads (DP≤4) were considered as missing values.

#### Filtering and missing value imputing

For every polymorphic site between the parental strains, we compared the polymorphisms in the segregants and the parental strains to infer which allele was inherited. We further excluded polymorphisms (i) that could not be called (or correctly called) in the parental strains based on RNA-seq data, (ii) that could be called in less than 70% of the segregants, and (iii) with a lower allele frequency of less than 20%. As previously discussed^18^, genotypes called differing from the two direct flanking markers in more than one segregant (93 cases) probably denote erroneous genotype calls; the corresponding polymorphisms were excluded from the analysis. This resulted in 25,590 polymorphisms that were considered as genetic markers.

Finally missing genotype values were inferred from the two neighboring polymorphisms if those were each within 20 kb of the polymorphism of interest, and were inherited from the same parental strain^18^ (Supplementary Data S3).

#### Assembling a set of markers for linkage analyses

Adjacent markers with the same segregation pattern across all segregants were collapsed into one unique marker, resulting in a set of 3,593 unique mapping genotypic markers (Supplementary Data S13). Thus, each marker represents a genomic interval in which all polymorphisms are in full linkage disequilibrium in the cross.

### Gene expression quantification and normalization

In previous work, we have shown that accounting for individual genome variations for RNA-seq alignment improved gene expression quantification and deflated the number of falsely detected local eQTLs^18^. Therefore we used the strategy we had previously developed to map RNA-seq reads. It consists of generating a strain-specific genome, for each segregant, against which the corresponding reads are aligned.

First, we generated both strain-specific genome sequences and strain-specific annotations from the reference genome sequence (SaCer3), the reference genome annotation and the genomic variations information (VCF files) previously created using RNA-seq data. Then RNA-seq reads were aligned to the corresponding strain specific genomes using STAR (ver 2.5.0a, ^40^) with the following options: `--alignIntronMin 10 --quantMode GeneCounts`. The gene-specific read counts (strand specific) generated by STAR were used to quantify gene expression.

RNA-seq coverage was computed by dividing the sum of the length of all reads by the sum of the length the coding regions of the quantified transcripts per sample.

Raw read counts were normalized using *rlog* method of DESeq2 ^41^. Normalized data were further corrected for effects due to culture batches using the non-parametric empirical Bayes framework ComBat^42^ (Supplementary Data S2). Normalized and batch corrected read counts were corrected for gene length as follows for gene *i* in sample *j*:

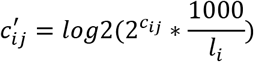

where *l*_*i*_ is the length of the coding region of gene *i*, excluding intronic regions.

### Proteomics

#### Sample preparation and phospho enrichment

Cell pellets were resuspended in lysis buffer containing 8 M urea, 0.1 M NH4HCO3, and 5 mM EDTA and cells were disrupted by glass bead beating (5 times for 5 min at 4°C, allowing the samples to cool down between cycles). The total protein amount from the pooled supernatants was determined by BCA Protein Assay Kit (Thermo, USA). Three milligrams of extracted yeast proteins were reduced with 5 mM TCEP at 37 °C for 30 min and alkylated with 12 mM iodoacetamide at room temperature in the dark for 30 min. The samples were then diluted with 0.1 M NH_4_HCO_3_ to a final concentration of 1 M urea and the proteins were digested with sequencing-grade porcine trypsin (Promega, Switzerland) at a final enzyme:substrate ratio of 1:100 (w/w). Digestion was stopped by adding formic acid to a final concentration of 1%. Peptide mixtures were desalted using 3cc reverse phase cartridges (Sep-Pak tC18, Waters, USA) and according to the following procedure: washing of column with one volume of 100% methanol, washing with one volume of 50% acetonitrile, washing with 3 volumes of 0.1% formic acid, loading acidified sample, reloading flow-through, washing column with sample with 3 volumes of 0.1% formic acid, and eluting sample with two volumes of 50% acetonitrile in 0.1% formic acid. Peptides were dried using a vacuum centrifuge and resolubilized in 100 μl of 0.1% formic acid. Retention time standard peptides (iRT-Kit, Biognosys, Switzerland) were spiked into the samples before they were analyzed by LC-MS for total protein abundances (“non-enriched samples”). The remaining 95 μl were supplemented with 300 μl of an overnight re-crystalized and cleared up phthalic acid solution prepared by carefully dissolving 5 g of phthalic acid in 50 ml of 80% acetonitrile before adding 1.75 ml of trifluoroacetic acid. The samples were then enriched for phosphopeptides by incubating for 1 hour under rotation with 1.25 mg of TiO2 resin (GLscience, Japan) pre-equilibrated twice with 500 μl of methanol, and twice with 500 μl of phthalic acid solution. Peptides bound to the TiO2 resin were then washed twice with 500 μl phthalic acid solution, then twice with 80% acetonitrile with 0.1% formic acid, and finally twice with 0.1% formic acid. The phosphopeptides were eluted from the beads twice with 150 μl of 0.3 M ammonium hydroxide at pH 10.5 and immediately acidified again with 50ul of 5% trifluoroacetic acid to reach about pH 2.0. The enriched phosphopeptides were desalted on microspin columns (The Nest Group, USA) with the protocol described above, dried using a vacuum centrifuge, and resolubilized in 10 μl of 0.1% formic acid. Again, retention time standard peptides (iRT-Kit, Biognosys, Switzerland) were spiked into the samples before they were analyzed by LC-MS for total peptide abundances (“phosphopeptides samples”).

#### LC-MS data acquisition

The peptide concentration in all samples was measured on a NanoDrop at OD280 and normalized to allow injection ~1 μg of material into the mass spectrometer. The samples were randomized and then either injected individually for SWATH-MS acquisition or pooled and injected in technical duplicates for shotgun acquisition. The LC-MS acquisitions were performed on an AB Sciex 5600 TripleTOF coupled to a NanoLC2Dplus HPLC system. The liquid chromatographic separation and mass spectrometric acquisition parameters were essentially as described earlier^43^. The peptide separation was performed on a 75 μm diameter PicoTip/PicoFrit emitter packed with 20 cm of Magic C18 AQ 3 resin using a 2-35% buffer B at 300 nl/min (buffer A: 2% acetonitrile, 0.1% formic acid; buffer B: 98% acetonitrile, 0.1% formic acid). For shotgun experiments, the mass spectrometer was operated with a “top 20” method, with a 500-ms survey scan followed by a maximum of 20 MS/MS events of 150 ms each. The MS/MS selection was set for precursors exceeding 200 counts per second and charge states greater than 2. The selected precursors were then added to a dynamic exclusion list for 20 s. Ions were isolated using a quadrupole resolution of 0.7 amu and fragmented in the collision cell using the collision energy equation

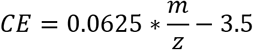

with a collision energy spread of 15 eV. For SWATH-MS acquisition, a 100-ms survey scan was followed by a series of 32 consecutive MS/MS events of 100 ms each with 25 amu precursor isolation with 1 amu overlap. The sequential precursor isolation window set-up was as follows: 400–425, 424–450, 449–475,…, 1174–1200 m/z. The collision energy for each window was determined based on the collision energy for a putative doubly charged ion centered in the respective window using the same equation as above with a collision energy spread of 15 eV.

All the MS data files were visually inspected and curated at this stage for low total ion chromatogram intensities, and the corresponding samples re-injected, if possible. This resulted in a final set of 179 SWATH data files for non-enriched and 179 SWATH data files for phospho-enriched samples (Supplementary Data S1) that were used for data extraction. Similarly, 40 DDA files for non-enriched and 30 DDA files for phospho-enriched samples were selected for database searching and library generation.

#### LC-MS database searching

The shotgun data was searched with Sorcerer-Sequest (TurboSequest v4.0.3rev11 running on a Sage-N Sorcerer v4.0.4) and Mascot (version 2.3.0) against the SGD database (release 03 Feb. 2011, containing 6,750 yeast protein entries, concatenated with 6,750 corresponding “tryptic peptide pseudo-reverse” decoy protein sequences). For the search, we allowed for semi-tryptic peptides and up to two missed cleavages per peptide. For the non-enriched samples, we used carbamidomethylation as a fixed modification on cysteine residues and oxidation as variable modification on methionine residues. For the phospho-enriched samples, we additionally allowed for phosphoralytion as variable modification on serine, threonine, and tyrosine residues. The Sequest and Mascot search results were converted to pep.xml and then combined using iProphet (included in TPP version 4.5.2) both for the non-enriched and for the phospho-enriched samples. Both search results were filtered at 1% FDR by decoy counting at the peptide spectrum matches (PSM) level, resulting in a total of 698,652 identified spectra, 26,893 unique peptides, and 4,310 proteins for the non-enriched sample set; in the phospho-enriched sample set there were a total of 224,551 identified spectra, 16,515 unique peptides (thereof 14,466 unique phosphopeptides), and 2,333 proteins (thereof 1,911 phospho-proteins). Those data were compiled into two spectra libraries (one ‘non-enriched’ and one ‘phospho-enriched’) using SpectraST (included in TPP 4.5.2) essentially as described earlier^44^, including the specific splitting of the consensus spectra when MS/MS scans identifying the same peptide sequence were recorded more than 2 minutes apart, also described earlier^44^. Those ‘split peptide assays’ were given different protein entry names labeled Subgroup_0_ProteinX to Subgroup_N_ProteinX, respectively. The fragment ion coordinates for the peptides contained the top 6 most intense (singly or doubly charged) y or b fragment ions for each spectrum, excluding those in the SWATH precursor isolation window for the corresponding peptide. The non-enriched assay library comprised assays for 19,473 peptides (thereof 18,074 proteotypic peptides matching a total of 3,119 unique proteins). The phospho-enriched assay library comprised assays for 14,339 peptides (thereof 12,969 phosphopeptide sequences) or assays for 13,786 proteotypic peptides (thereof 12,678 proteotypic phosphopeptides, matching a total of 1,676 unique phosphoproteins).

The SWATH-MS data extraction was performed using the iPortal workflow manager^45^ calling OpenSWATH (openMS v. 1.10) ^46^ and pyProphet ^47^. The precursors were then re-aligned across runs using TRIC ^48^. The two resulting SWATH identification result files contained a total of 18,273 identified peptides (thereof 16,922 proteotypic peptides matching a total of 2,940 proteins) for the non-enriched datasets; in the phospho-enriched datasets there were 13,748 identified peptides (thereof 12,412 phosphopeptides) or 13,218 proteotypic peptides (thereof 12,139 proteotypic phosphopeptides matching a total of 2,247 unique phosphoproteins). After alignment, we used a set of in-house scripts to compare the chromatographic elution profiles of the various isobaric phosphopeptide isoforms matching a same delocalized peptide form (peptide sequence + number of phosphorylations) within each single run and to eventually group those co-eluting phosphopeptide assays into the proper corresponding number of phospho-peak clusters (labeled _cluster0 to _clusterN, respectively). The phospho-peak clusters were then consistently re-numbered across runs and those were used as input to mapDIA^49^ to select for the best suitable transitions and peptides for quantification. This resulted in the final peptide and protein quantification matrices for the non-enriched and phospho-enriched datasets that were used for further processing.

#### Preprocessing of proteome data

First, features detected after 7000 s and those corresponding to decoys, reverse proteins, or not unique peptides were removed. We also removed fragments with oxidized methionine and their corresponding non-oxidized fragments. Next, native retention times were converted to iRTs^51^. Fragments corresponding to peptides, whose sequence was existing only in the reference proteome (i.e. BY1416 background) and not in RM11-1a background were excluded from subsequent analysis. Normalization of the fragment-level data and aggregation into peptides and protein-level data was performed using mapDIA^49^ with the following options: `NORMALIZATION = RT 10, MIN_CORREL = 0.3, MIN_FRAG_PER_PEP = 2, MIN_PEP_PER_PROT = 1` and a maximum of 20% missing data for each fragment. Finally, abundance data were further corrected for effects due to culture batches and proteomics measurement batches using the non-parametric empirical Bayes framework ComBat^42^.

#### Preprocessing of phospho-proteome data

First fragments detected after 6000 sec, corresponding to non-phosphorylated peptides (0P), corresponding to non-unique proteins, and oxidized were removed. As for non-phosphorylated proteomics data, native retention times were replaced by iRT^51^ and fragments corresponding to peptides, which sequence was not existing in RM11-1a background were excluded from subsequent analysis. To normalize the fragment level data and to aggregate them at the phosphopeptide cluster level we used mapDIA^49^. Fragments derived from phosphopeptides belonging to the same phosphopeptides cluster were aggregated together. The following mapDIA options were used: `NORMALIZATION = RT 10, MIN_CORREL = 0.3, MIN_FRAG_PER_PEP = 2` and a maximum proportion of missing data of 20% for each fragment as for the non-phospho proteomics data. Data were corrected for culture batch and well as phospho-proteomics measurements batch effects using ComBat^42^.

### Derivation of the post-transcriptional traits (protein abundance regressed to RNA levels) and phosphorylation levels (phosphopeptide abundance regressed to protein abundance)

In order (i) to separate the changes in protein abundance due to RNA changes from the post-transcriptional specific regulation, and (ii) to distinguish changes in phosphopeptide abundance due to protein abundance changes from modification of the phosphorylation levels, we have generated regressed traits (as already proposed in ^8^). The same procedure has been applied for both traits and will be detailed below using the phosphopeptide abundance example.

First, for each pair of corresponding traits (i.e. a phosphopeptide and its protein of origin) relative abundances across sample were normalized using a modified transformation to standard score (i.e. centering and scaling). To compute mean and standard deviation for this normalization, only the values corresponding to the samples, in which measurements were available for both phosphopeptides and protein were used. Then, for each pair, normalized phosphopeptide data were regressed on protein data using a robust linear regression using a MM-estimate^52^, initialized by an S-estimate using Hubber’s weights function and using an M-estimator as final estimate using a Tukey’s biweight function as implemented MM-estimation option in the *rlm* function of the MASS R package^53^. The residuals of these regression were then used to as trait in the subsequences analyses.

### QTL mapping

We employed a previously developed QTL detection method based on Random Forest (RF), that was further modified to improve its performance ^18,21^.

To correct for population substructure we include population structure as a covariate in the model, as we previously proposed^21^, but in a slightly different manner. First, the genotype matrix was normalized as described in ^54^ (see equations (1) to (3) there). Then we carried out a singular value decomposition on the normalized genotype matrix followed by an eigenvector decomposition. We then selected those eigenvectors corresponding to the top seven eigenvalues as covariates for the QTL mapping (additional predictors for growing the Random Forests). These first seven vectors explained more than 25% of the genotype variance (Supplementary Data S14).

We adapted the approach described in ^18^ by using a different score describing the importance of each predictor in the Random Forest (variable importance measure, VIM). While previous work relied on the selection frequency as the VIM, i.e. the number of times a predictor was used in a forest, to quantify the importance of each predictor, we combined two previously published VIMs, RSS and PI^55^, to compute an informative combined score serving as the VIM in our study. While the RSS describes the average reduction in the sum of the squared residuals after splitting a node with a predictor, the PI refers to the relative reduction in predictive accuracy after permuting the information for the respective predictor.

RSS and PI are combined in the following way to generate a more robust score *S_i_* for predictor *i*:

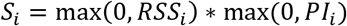

We averaged replicates for the same strain to avoid the detection of false positive QTL. Consecutive markers linked to the same trait and/or markers in high LD (Pearson’s *r* > 0.8) linked to the same trait were counted as a single linkage as we previously described^18^.

### QTL hotspot detection

To formally identify hotspots in our dataset, the genome was divided into 40-kb bins (293 bins, bins at chromosome extremities could be bigger). As proposed before^15^, if the linkages were randomly distributed across the genome, the number of linkages in each bin would be expected to follow a Poisson distribution with the mean of N_*linkage*_/N_*bin*_. We use this distribution to estimate the highest number of linkages that a bin could contain at a probability lower than 0.01. These numbers were 30 for eQTL, 13 for pQTL, 9 for ptQTL, 11 for phQTL, and 4 for phResQTL. Bins with a sufficient number of QTLs at a given molecular layer were considered as QTL-hotspots. Consecutive bins on the same chromosome with enough QTLs were combined to single hotspots.

### Integration of growth QTL

QTL affecting growth under various conditions were taken from ^20^. If the reported peak position of the growth QTL was within 50 kb of the middle of the positions of the loci that affected the most traits on each of the molecular layers, we considered them to be the same QTL.

### Identification of local QTL

QTLs were considered to be local to their target if the QTL contained at least one genetic marker that had a correlation of 0.8 or higher with one of the markers that are directly up- or downstream of the target gene (Pearson’s *r*). For eQTL, the target gene corresponds to the gene, whose expression is investigated; for pQTL it corresponds to the gene encoding the protein, whose abundance is studied; for phQTL and phResQTL it corresponds to the gene encoding the protein of which the phosphopeptide is part of.

### GO enrichment

Gene Ontology (GO) enrichment analyses were performed using topGO. We performed Fisher’s exact tests with the “*weight01*” algorithm and a minimal nodesize of 10 ^56^. We used annotations from SacCer3.

### Broad-sense heritability estimation

Broad-sense heritability estimates were computed based on replicate measurements of some strains, as described elsewhere^20^. We used six different segregants with three replicates each, as well as the parents with six (BY) and eight (RM) replicates each. In short, the ‘lmer’ function from the lme4 R package was used to create a linear mixed-effects model with the phenotype as the response and the segregant labels as random effects57. The variance components *σ*^2^_*G*_ (the variance due to genetic effects, i.e., different segregants), and *σ*^2^_*E*_, (the error variance), were extracted, and broad-sense heritability was calculated as H^2^ = *σ*^2^_*G*_/ (*σ*^2^_*G*_+*σ*^2^_*E*_). Standard errors were calculated using the delete-one jackknife procedure, as proposed previously^20^. Random distributions of heritability estimates for each molecular layer were generated by permuting the strain labels and compared to the real distributions using a Mann-Whitney test. In order to be able to compare heritability estimates between the different molecular levels, we restricted the analysis to the set of 402 proteins where we had measurements for expression, protein, and (at least one) phosphopeptide.

### Polymorphisms in and around genes

To investigate the cause of local QTL, we counted polymorphisms in the upstream region, downstream region, 3’ and 5’ UTRs, coding sequence and amino acid sequence of each coding gene between the BY and RM genomes. We considered all SNPs reported by Bloom et al. ^20^. We excluded all genes with insertions and deletions from this analysis. UTR-annotations were downloaded from www.yeastgenome.org in October 2017. If the UTR was reported multiple times with differing lengths, we used the largest annotation. The up- and downstream regions of a gene spanned 2kb each and began at the outer borders of the UTRs relative to the gene of interest. The number of coding polymorphisms was defined as the number of amino acid changes in a protein between BY and RM. Multiple SNPs in the same codon were only counted once.

Figure 3c shows only genes with available UTR annotations and available measurements on the protein level. Genes with indels are not included.

For each phosphosite we identified the closest polymorphism in the amino acid sequence space by computing the absolute difference of the position of the modified serine and all polymorphic amino acids within the protein and using the minimum of those distances. Note that phosphopeptides were only included in this study if they didn’t have any polymorphisms.

### Annotation of protein complexes

We used previously published annotations for protein complexes (ftp://ftp.ebi.ac.uk/pub/databases/intact/complex/current/complextab/saccharomyces_cerevisiae.tsv, ^58^). The annotated complex members included proteins and small molecules such as ions. We only considered those proteins as members of complexes that shared a complex with at least one other protein.

To investigate the influence of the stoichiometry of protein complexes on protein levels, we computed the correlation between levels of proteins of the same complex. Here we only considered genes (i) for which protein and transcript levels were available, and (ii) which were annotated for complexes that did not have any overlap with other complexes. Nucleolar and ribosomal proteins were excluded from this analysis, since we already showed that they were enriched in genes with buffered protein levels (‘buffered’ in Figure 3b). Taking them into account might have biased the results from the analysis of complex protein pairs. In total we considered 286 unique pairs of proteins that each shared a complex. These pairs corresponded to 188 genes.

We then used Fisher’s exact tests to determine if pairs of genes that share a complex were affected by the same QTL at a significantly different frequency than pairs of genes that do not share a complex. Traits were considered to be affected by the same QTL if any marker was significant for both traits.

### Classification of QTL based on their effects on the transcriptome and proteome

To analyze how QTL affect the proteome, we classified eQTL into three distinct classes: (i) eQTL with ‘similar’ effects on the transcriptome and proteome, (ii) ‘buffered’ eQTL, and (iii) ‘enhanced’ eQTL. We computed the effect *E* of a QTL at locus *i* on a molecular trait *j* as a log2 fold change by subtracting the average of the log2-transformed trait values *t* of all strains with the RM-allele at this locus from the average trait value for all strains with BY-allele.

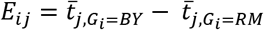

The eQTLs were classified based on the difference of their effects on the target transcript and the encoded protein:

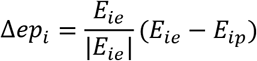

where *E*_*ie*_ and *E*_*ip*_ are the effects of eQTL *i* on the mRNA and protein levels, respectively. If the absolute difference |Δ*ep*_*i*_| was below 0.15 the eQTL effect was classified as ‘similar’. If Δ*ep*_*i*_ was above +0.15, it was classified as ‘buffered’ and if the effect difference was below −0.15 (i.e. Δ*ep*_*i*_ < −0.15) it was classified as ‘enhanced’.

ptQTL at FDR<10% that didn’t overlap with an eQTL for the same gene were added as a fourth class (‘protein only’).

The effects of local and distant eQTL on the proteome was compared, by generating simple least squares regression models for local and distant eQTL separately. Here we considered the log2 fold change on the protein level as the dependent variable and the log2 fold change on the transcript level as the independent variable. Models were generated with the *lm* function of the *stats* package in R^59^.

### Enrichment of targets of kinases and phosphatases among HAP1-targets

We tested the phosphoproteins targeted by the *HAP1*-locus for enrichments in the previously annotated targets of a large number of kinases and phosphatases^34^. Here we only considered target proteins that were reported to be phosphorylated or dephosphorylated at serine residues. We also considered proteins that were measured in ^34^ but not found to be regulated by any genetic perturbation. Among the 315 proteins that were measured in both studies, a total of 45 proteins had at least one phResQTL at the *HAP1*-locus. We used one-sided Fisher’s exact tests to assess the significance of the enrichments of targets of a kinase or phosphatase among the genes that were also targeted by the *HAP1*-locus.

## Data accessibility

All raw MS data files and search results thereof have been deposited to the ProteomeXchange Consortium via the PRIDE partner repository with the dataset identifier PXD010893.

Raw RNA-seq reads for all 150 samples that were included in the eQTL-mapping were uploaded to ArrayExpress under the accession E-MTAB-8146.

## References

1. Emilsson V, Thorleifsson G, Zhang B, Leonardson AS, Zink F, Zhu J, Carlson S, Helgason A, Walters GB, Gunnarsdottir S, Mouy M, Steinthorsdottir V, Eiriksdottir GH, Bjornsdottir G, Reynisdottir I, Gudbjartsson D, Helgadottir A, Jonasdottir A, Jonasdottir A, Styrkarsdottir U, Gretarsdottir S, Magnusson KP, Stefansson H, Fossdal R, Kristjansson K, Gislason HG, Stefansson T, Leifsson BG, Thorsteinsdottir U, Lamb JR, Gulcher JR, Reitman ML, Kong A, Schadt EE, Stefansson K. Genetics of gene expression and its effect on disease. Nature (2008).

2. van der Sijde, M. R., Ng, A. & Fu, J. Systems genetics: From GWAS to disease pathways. Biochim. Biophys. Acta BBA - Mol. Basis Dis. 1842, 1903–1909 (2014).

3. Bali V, Bebok Z. Decoding mechanisms by which silent codon changes influence protein biogenesis and function. Int. J. Biochem. Cell Biol. (2015).

4. Henchoz S, Chi Y, Catarin B, Herskowitz I, Deshaies RJ, Peter M. Phosphorylation- and ubiquitin-dependent degradation of the cyclin-dependent kinase inhibitor Far1p in budding yeast. Genes Dev. (1997).

5. Ardito, F., Giuliani, M., Perrone, D., Troiano, G. & Muzio, L. L. The crucial role of protein phosphorylation in cell signaling and its use as targeted therapy (Review). Int. J. Mol. Med. 40, 271–280 (2017).

6. Miller, M. E. & Cross, F. R. Mechanisms Controlling Subcellular Localization of the G1 Cyclins Cln2p and Cln3p in Budding Yeast. Mol. Cell. Biol. 21, 6292–6311 (2001).

7. Albert FW, Bloom JS, Siegel J, Day L, Kruglyak L. Genetics of trans-regulatory variation in gene expression. eLife (2018).

8. Foss, E. J. et al. Genetic Variation Shapes Protein Networks Mainly through Non-transcriptional Mechanisms. PLoS Biol. 9, e1001144 (2011).

9. Holdt, L. M. et al. Quantitative Trait Loci Mapping of the Mouse Plasma Proteome (pQTL). Genetics 193, 601–608 (2013).

10. Wu, L. et al. Variation and genetic control of protein abundance in humans. Nature 499, 79–82 (2013).

11. Picotti, P. et al. A complete mass-spectrometric map of the yeast proteome applied to quantitative trait analysis. Nature 494, 266–270 (2013).

12. Gowda GA, Djukovic D. Overview of mass spectrometry-based metabolomics: opportunities and challenges. Methods Mol. Biol. Clifton NJ (2015).

13. Chick, J. M. et al. Defining the consequences of genetic variation on a proteome-wide scale. Nature 534, 500–505 (2016).

14. Civelek, M. & Lusis, A. J. Systems genetics approaches to understand complex traits. Nat. Rev. Genet. 15, 34–48 (2014).

15. Brem, R., Yvert, G., Clinton, R. & Kruglyak, L. Genetic Dissection of Transcriptional Regulation in. Nature 411, 41 (2002).

16. Foss, E. J. et al. Genetic basis of proteome variation in yeast. Nat. Genet. 39, 1369–1375 (2007).

17. Albert, F. W., Treusch, S., Shockley, A. H., Bloom, J. S. & Kruglyak, L. Genetics of single-cell protein abundance variation in large yeast populations. Nature 506, 494–497 (2014).

18. Clement-Ziza, M. et al. Natural genetic variation impacts expression levels of coding, non-coding, and antisense transcripts in fission yeast. Mol. Syst. Biol. 10, 764–764 (2014).

19. Brem RB, Kruglyak L. The landscape of genetic complexity across 5,700 gene expression traits in yeast. Proc. Natl. Acad. Sci. U. S. A. (2005).

20. Bloom, J. S., Ehrenreich, I. M., Loo, W. T., Lite, T.-L. V. & Kruglyak, L. Finding the sources of missing heritability in a yeast cross. Nature 494, 234–237 (2013).

21. Michaelson, J. J., Alberts, R., Schughart, K. & Beyer, A. Data-driven assessment of eQTL mapping methods. BMC Genomics 11, 502 (2010).

22. Smith, E. N. & Kruglyak, L. Gene–Environment Interaction in Yeast Gene Expression. PLoS Biol. 6, e83 (2008).

23. Yvert, G. et al. Trans-acting regulatory variation in Saccharomyces cerevisiae and the role of transcription factors. Nat. Genet. 35, 57–64 (2003).

24. Tsay, Y. F., Thompson, J. R., Rotenberg, M. O., Larkin, J. C. & Woolford, J. L. Ribosomal protein synthesis is not regulated at the translational level in Saccharomyces cerevisiae: balanced accumulation of ribosomal proteins L16 and rp59 is mediated by turnover of excess protein. Genes Dev. (1988). doi:10.1101/gad.2.6.664

25. Liu Y, Mi Y, Mueller T, Kreibich S, Williams EG, Van Drogen A, Borel C, Frank M, Germain PL, Bludau I, Mehnert M, Seifert M, Emmenlauer M, Sorg I, Bezrukov F, Bena FS, Zhou H, Dehio C, Testa G, Saez-Rodriguez J, Antonarakis SE, Hardt WD, Aebersold R. Multi-omic measurements of heterogeneity in HeLa cells across laboratories. Nat. Biotechnol. (2019).

26. Liu Y, Borel C, Li L, Müller T, Williams EG, Germain PL, Buljan M, Sajic T, Boersema PJ, Shao W, Faini M, Testa G, Beyer A, Antonarakis SE, Aebersold R. Systematic proteome and proteostasis profiling in human Trisomy 21 fibroblast cells. Nat. Commun. (2018).

27. Jüschke C, Dohnal I, Pichler P, Harzer H, Swart R, Ammerer G, Mechtler K, Knoblich JA. Transcriptome and proteome quantification of a tumor model provides novel insights into post-transcriptional gene regulation. Genome Biol. (2014).

28. Garcia-Barrio M, Dong J, Ufano S, Hinnebusch AG. Association of GCN1-GCN20 regulatory complex with the N-terminus of eIF2alpha kinase GCN2 is required for GCN2 activation. EMBO J. (2000).

29. Ben-Levy R, Leighton IA, Doza YN, Attwood P, Morrice N, Marshall CJ, Cohen P. Identification of novel phosphorylation sites required for activation of MAPKAP kinase-2. EMBO J. (1996).

30. Tatchell, K., Robinson, L. C. & Breitenbach, M. RAS2 of Saccharomyces cerevisiae is required for gluconeogenic growth and proper response to nutrient limitation. Proc. Natl. Acad. Sci. 82, 3785–3789 (1985).

31. Jian, D., Aili, Z., Xiaojia, B., Huansheng, Z. & Yun, H. Feedback regulation of Ras2 guanine nucleotide exchange factor (Ras2-GEF) activity of Cdc25p by Cdc25p phosphorylation in the yeast Saccharomyces cerevisiae. FEBS Lett. (2010). doi:10.1016/j.febslet.2010.11.006

32. Verdière J, Creusot F, Guérineau M. Regulation of the expression of iso 2-cytochrome c gene in S. cerevisiae: cloning of the positive regulatory gene CYP1 and identification of the region of its target sequence on the structural gene CYP3. Mol. Gen. Genet. MGG (1985).

33. Rutter J, Probst BL, McKnight SL. Coordinate regulation of sugar flux and translation by PAS kinase. Cell (2002).

34. Bodenmiller B, Wanka S, Kraft C, Urban J, Campbell D, Pedrioli PG, Gerrits B, Picotti P, Lam H, Vitek O, Brusniak MY, Roschitzki B, Zhang C, Shokat KM, Schlapbach R, Colman-Lerner A, Nolan GP, Nesvizhskii AI, Peter M, Loewith R, von Mering C, Aebersold R. Phosphoproteomic analysis reveals interconnected system-wide responses to perturbations of kinases and phosphatases in yeast. Sci. Signal. (2010).

35. Klipp, E. & Liebermeister, W. Mathematical modeling of intracellular signaling pathways. BMC Neurosci. 7, S10 (2006).

36. Mas, G. et al. Recruitment of a chromatin remodelling complex by the Hog1 MAP kinase to stress genes. EMBO J. 28, 326–336 (2009).

37. Bilsland-Marchesan, E., Ariño, J., Saito, H., Sunnerhagen, P. & Posas, F. Rck2 kinase is a substrate for the osmotic stress-activated mitogen-activated protein kinase Hog1. Mol. Cell. Biol. 20, 3887–3895 (2000).

38. Kim D, Pertea G, Trapnell C, Pimentel H, Kelley R, Salzberg SL. TopHat2: accurate alignment of transcriptomes in the presence of insertions, deletions and gene fusions. Genome Biol. (2014).

39. Van der Auwera GA, Carneiro MO, Hartl C, Poplin R, Del Angel G, Levy-Moonshine A, Jordan T, Shakir K, Roazen D, Thibault J, Banks E, Garimella KV, Altshuler D, Gabriel S, DePristo MA. From FastQ data to high confidence variant calls: the Genome Analysis Toolkit best practices pipeline. Curr. Protoc. Bioinforma. (2016).

40. Dobin A, Davis CA, Schlesinger F, Drenkow J, Zaleski C, Jha S, Batut P, Chaisson M, Gingeras TR. STAR: ultrafast universal RNA-seq aligner. Bioinforma. Oxf. Engl. (2012).

41. Love MI, Huber W, Anders S. Moderated estimation of fold change and dispersion for RNA-seq data with DESeq2. Genome Biol. (2015).

42. Johnson WE, Li C, Rabinovic A. Adjusting batch effects in microarray expression data using empirical Bayes methods. Biostat. Oxf. Engl. (2007).

43. Selevsek N, Chang CY, Gillet LC, Navarro P, Bernhardt OM, Reiter L, Cheng LY, Vitek O, Aebersold R. Reproducible and consistent quantification of the Saccharomyces cerevisiae proteome by SWATH-mass spectrometry. Mol. Cell. Proteomics MCP (2015).

44. Schubert OT, Gillet LC, Collins BC, Navarro P, Rosenberger G, Wolski WE, Lam H, Amodei D, Mallick P, MacLean B, Aebersold R. Building high-quality assay libraries for targeted analysis of SWATH MS data. Nat. Protoc. (2015).

45. Kunszt, P. et al. iPortal: the swiss grid proteomics portal: Requirements and new features based on experience and usability considerations. Concurr. Comput. Pract. Exp. 27, 433–445 (2015).

46. Röst HL, Rosenberger G, Navarro P, Gillet L, Miladinović SM, Schubert OT, Wolski W, Collins BC, Malmström J, Malmström L, Aebersold R. OpenSWATH enables automated, targeted analysis of data-independent acquisition MS data. Nat. Biotechnol. (2014).

47. Teleman J, Röst HL, Rosenberger G, Schmitt U, Malmström L, Malmström J, Levander F. DIANA–algorithmic improvements for analysis of data-independent acquisition MS data. Bioinforma. Oxf. Engl. (2015).

48. Röst HL, Liu Y, D’Agostino G, Zanella M, Navarro P, Rosenberger G, Collins BC, Gillet L, Testa G, Malmström L, Aebersold R. TRIC: an automated alignment strategy for reproducible protein quantification in targeted proteomics. Nat. Methods (2017).

49. Teo G, Kim S, Tsou CC, Collins B, Gingras AC, Nesvizhskii AI, Choi H. mapDIA: Preprocessing and statistical analysis of quantitative proteomics data from data independent acquisition mass spectrometry. J. Proteomics (2015).

50. Vizcaíno JA, Côté RG, Csordas A, Dianes JA, Fabregat A, Foster JM, Griss J, Alpi E, Birim M, Contell J, O’Kelly G, Schoenegger A, Ovelleiro D, Pérez-Riverol Y, Reisinger F, Ríos D, Wang R, Hermjakob H. The PRoteomics IDEntifications (PRIDE) database and associated tools: status in 2013. Nucleic Acids Res. (2012).

51. Escher C, Reiter L, MacLean B, Ossola R, Herzog F, Chilton J, MacCoss MJ, Rinner O. Using iRT, a normalized retention time for more targeted measurement of peptides. Proteomics (2012).

52. Yohai, V. J. High Breakdown-Point and High Efficiency Robust Estimates for Regression. Ann. Stat. 15, 642–656 (1987).

53. Venables, W. N. & Ripley, B. D. Modern Applied Statistics with S. (Springer, 2002).

54. Patterson N, Price AL, Reich D. Population structure and eigenanalysis. PLoS Genet. (2007).

55. Liaw, A. & Wiener, M. Classification and regression by randomForest. R News 2, 18–22 (2002).

56. Alexa, A., Rahnenfuhrer, J. & Lengauer, T. Improved scoring of functional groups from gene expression data by decorrelating GO graph structure. Bioinformatics 22, 1600–1607 (2006).

57. Bates, D., Mächler, M., Bolker, B. & Walker, S. Fitting Linear Mixed-Effects Models Using lme4. J. Stat. Softw. 67, 1–48 (2015).

58. Meldal BH, Forner-Martinez O, Costanzo MC, Dana J, Demeter J, Dumousseau M, Dwight SS, Gaulton A, Licata L, Melidoni AN, Ricard-Blum S, Roechert B, Skyzypek MS, Tiwari M, Velankar S, Wong ED, Hermjakob H, Orchard S. The complex portal–an encyclopaedia of macromolecular complexes. Nucleic Acids Res. (2015).

59. R Core Team. R: A Language and Environment for Statistical Computing. (R Foundation for Statistical Computing, 2013).

