## Supplementary Materials for "Integration of transcriptome, proteome and phosphoproteome data elucidates the genetic control of molecular networks"

#### Supplementary Text

##### Supplementary Text S1

Chick *et al*. observed that, after correcting for transcript abundance, the significance of local pQTLs decreased more than that of distant pQTLs (Ref 13). Foss and colleagues detected markedly fewer local pQTLs than local eQTLs and concluded that protein levels are largely regulated in trans (Ref 8). However, the extent to which eQTLs impact protein levels might also depend on effect sizes of the eQTL. Indeed, earlier studies have established that the effects of local eQTLs are often larger than those of distant eQTLs (Ref 7). This was supported by our data: the average effect size of local eQTLs at FDR<10% was 26% larger than that of distant eQTLs.

Correspondingly, we found more local QTLs than expected by chance (p<2.2E-16 for all five types of molecular traits considered here, Fisher's exact test), which is also recognizable by the presence of diagonal bands in the QTL maps (Supplementary Figures S3-S7). Transcripts with a local eQTL had on average lower expression levels than those that were only affected by eQTLs acting in trans, further emphasizing that local QTLs are easier to detect (p<2.2E-16, Wilcoxon's rank sum test).

### Supplementary Figures

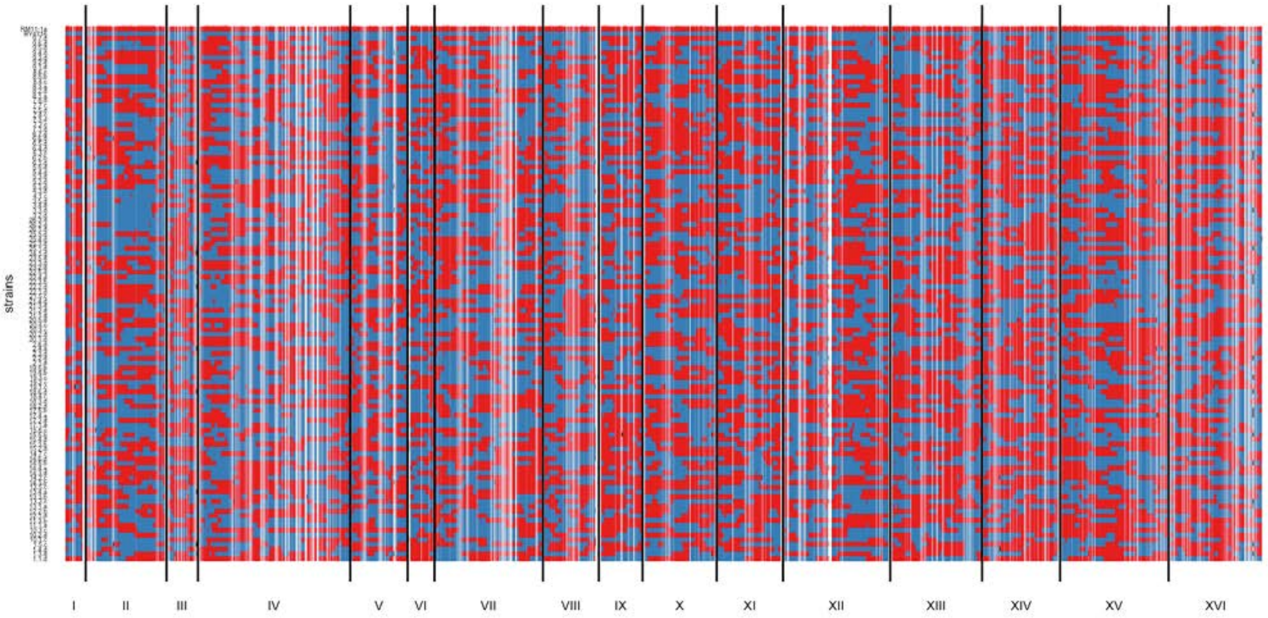


**Supplementary Figure S1** Genotypes of parental strains and segregants. Alleles called from variant positions in transcripts are shown in red and blue depending of the genome of origin. Alleles from RM are shown in red while alleles from BY are shown in blue.

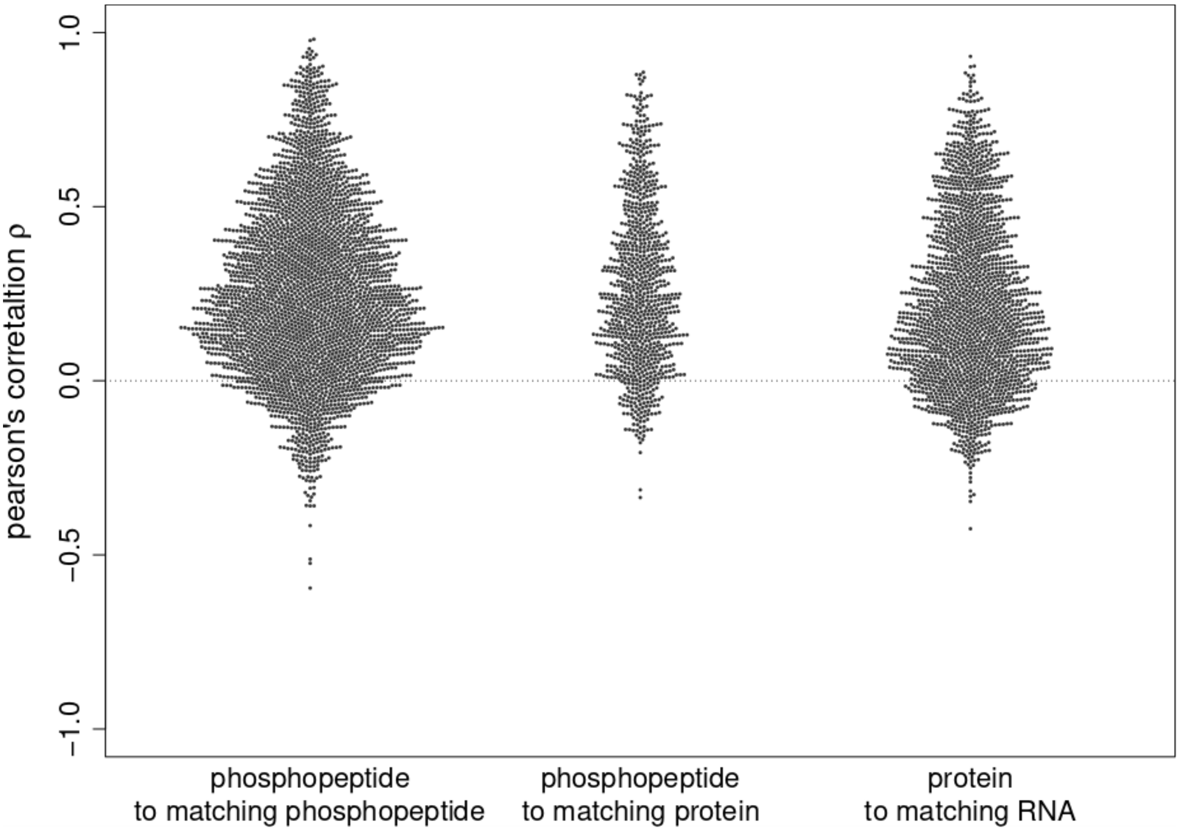


**Supplementary Figure S2** Correlation between molecular traits of the same gene. The pearson's correlations between phosphopeptides from the same protein (left), phosphopeptides and their host protein (middle) and that between protein levels and transcript levels of the same gene (right) are shown.

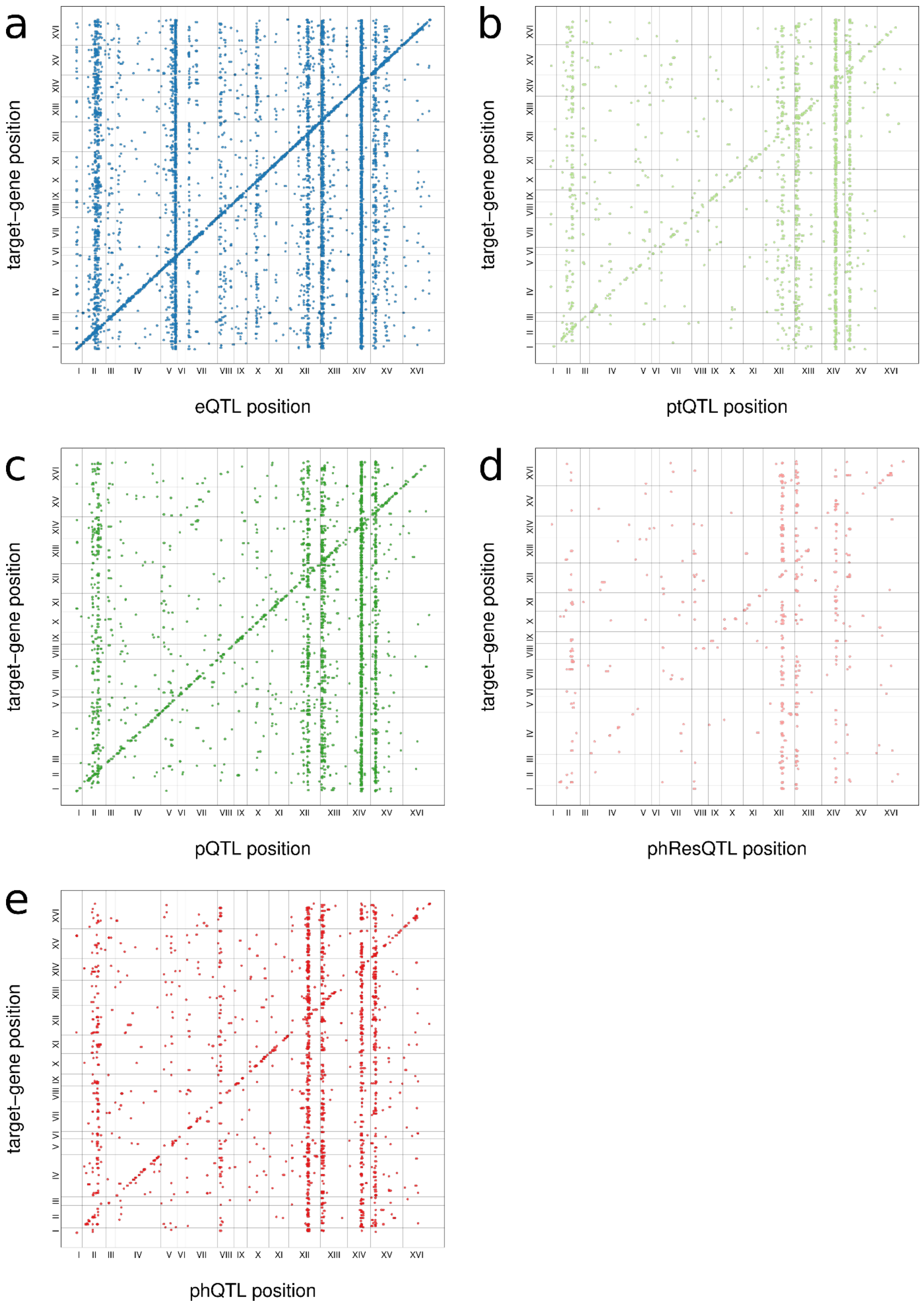


**Supplementary Figure S3** Significant QTL at FDR<10%. Associations between QTL and targets are shown as dots with X-coordinates showing the position of the QTL and Y-coordinates showing the position of the affected trait in the genome. Vertical bars indicate QTL-hotspots while the diagonal consists of local QTL. Shown are eQTL (**a**), ptQTL (**b**), pQTL (**c**), phResQTL (**d**), and phQTL (**e**).

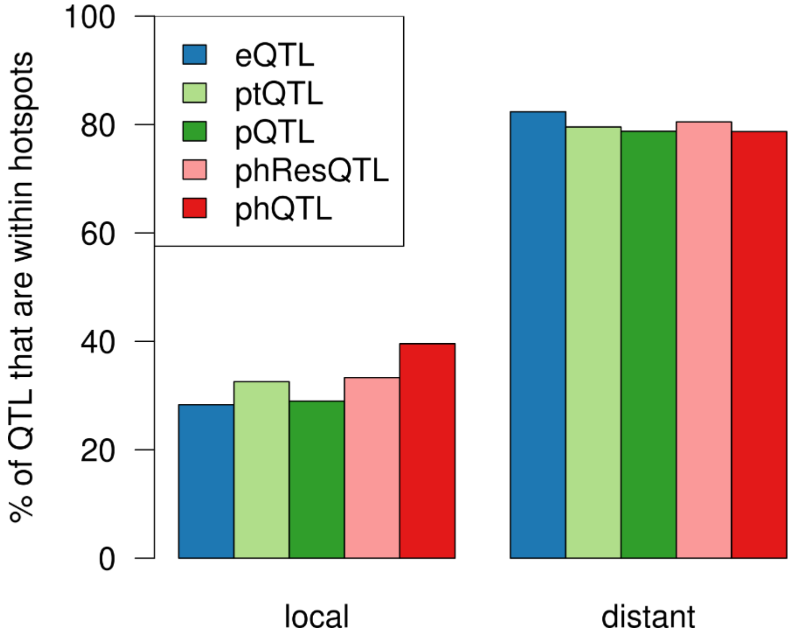


**Supplementary Figure S4** Proportion of local and distant QTL that are located in hotspots, show separately for each molecular layer.

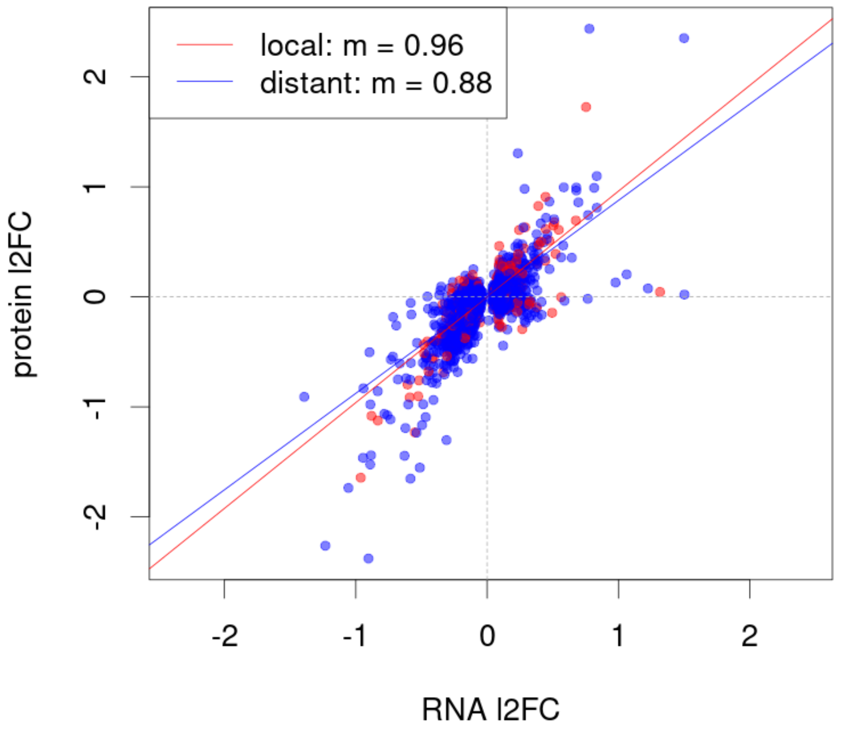


**Supplementary Figure S5** Transmission of local and distant eQTL effects to the proteome. For all significant eQTL at FDR<10% the effect on the transcript- and protein-levels of the target gene is shown. Local eQTL are shown as red dots, distant QTL are shown in blue. Protein-effects were regressed against transcripts-effects for each class of eQTL. Both linear models had similar.

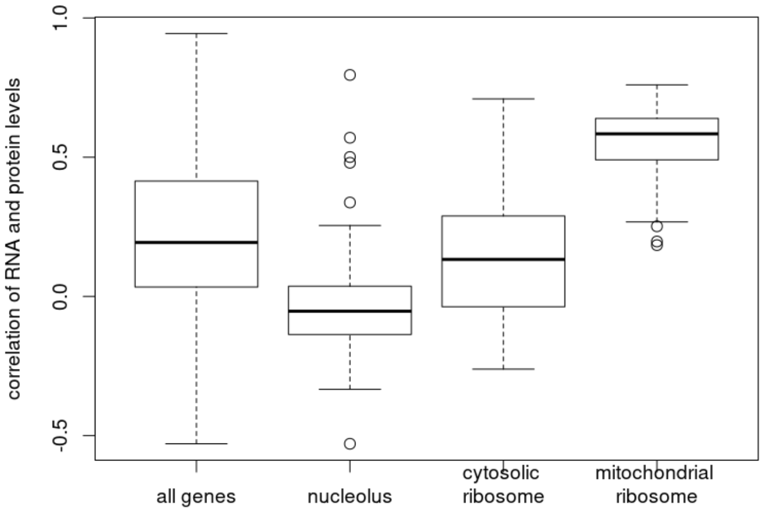


**Supplementary Figure S6** Correlation between transcript- and protein-levels for functional groups of genes. Levels of transcripts and proteins were correlated for all genes with available transcript- and protein-levels, genes annotated to the nucleolus (GO:0005730), the cytosolic ribosome (GO:0022626) and the mitochondrial ribosome (GO:0005761). Outlier are shown as circles outside of the whiskers of the boxplots.

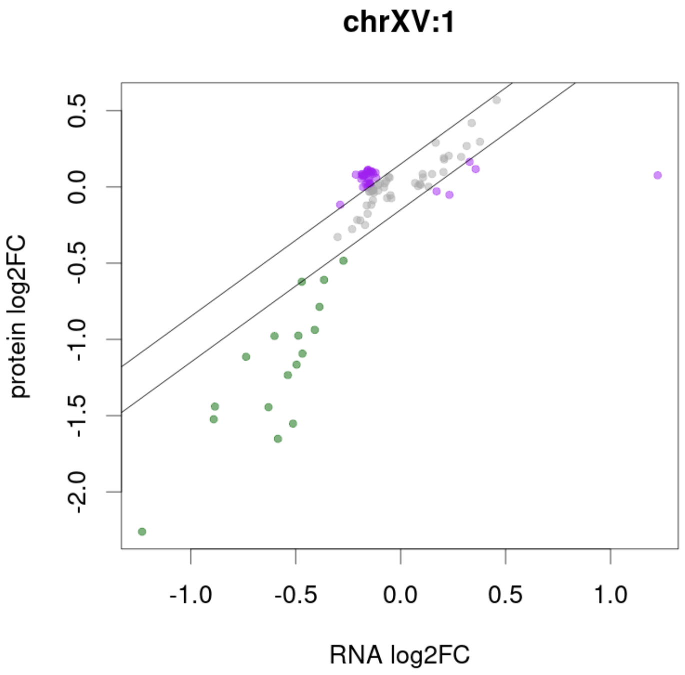


**Supplementary Figure S7** Transmission of eQTL effects from the IRA2 hotspot. Effects of eQTL at the IRA2-locus on the transcript- and protein-levels of the target genes at FDR<10%. Each dot represents a target gene and is colored according to the difference in its effect on transcript- and protein-levels as described in the main text.

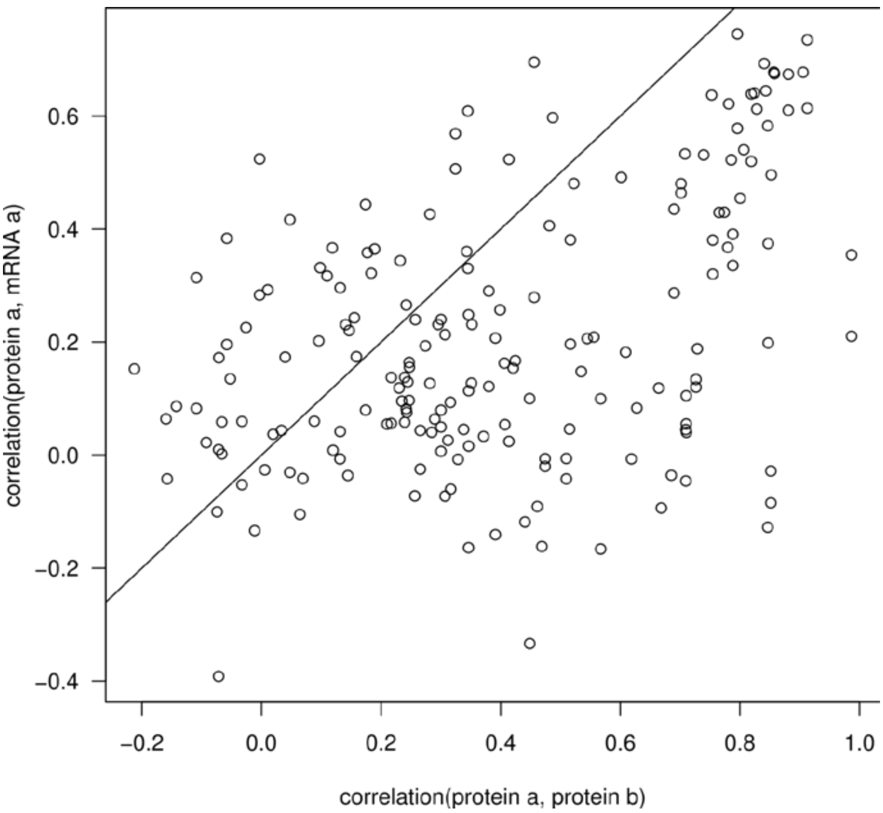


**Supplementary Figure S8** Correlation of protein levels of proteins in complexes to the levels of their own mRNA and to protein levels of the second protein of the same complex. Here we only considered complexes whose annotation did not overlap with other complexes and which had exactly two annotated members. Genes annotated to the nucleolus or as ribosomal proteins were not considered here.

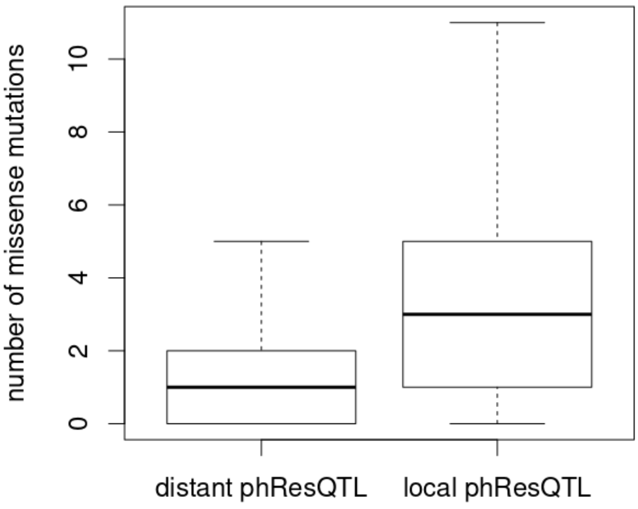


**Supplementary Figure S9** Number of missense mutations in proteins with phResQTL. The amount of missense mutations is shown for proteins with only distant phResQTL and for proteins with a local phResQTL.

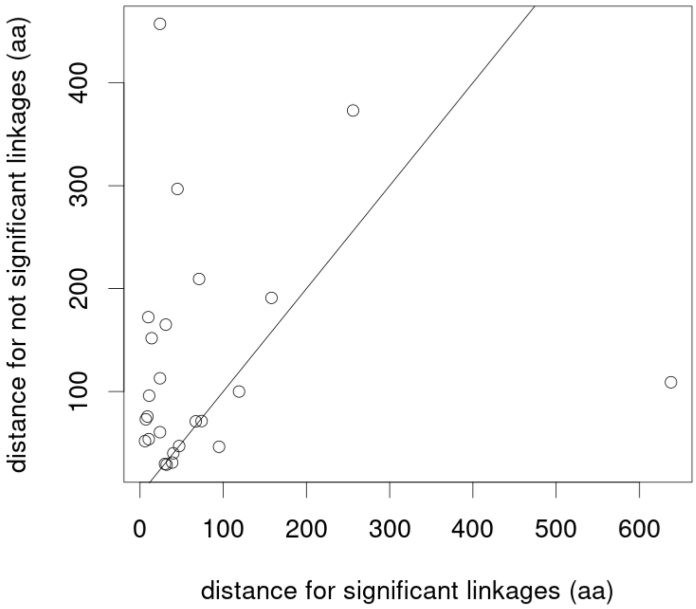


**Supplementary Figure S10** Distances to the closest missense mutations for multiple phosphosites on the same protein. For proteins with one phosphosite with a local phResQTL and a different phosphosite with a distant phResQTL the distance of both phosphosites to the closest missense mutation in the sequence space is shown. A solid black line shows the diagonal. Dots above the line represent proteins for which the phosphosite with the distant phResQTL is located further away from the closest missense mutation than the phosphosite with the local phResQTL. Dots under the line represent proteins where the reverse is true.

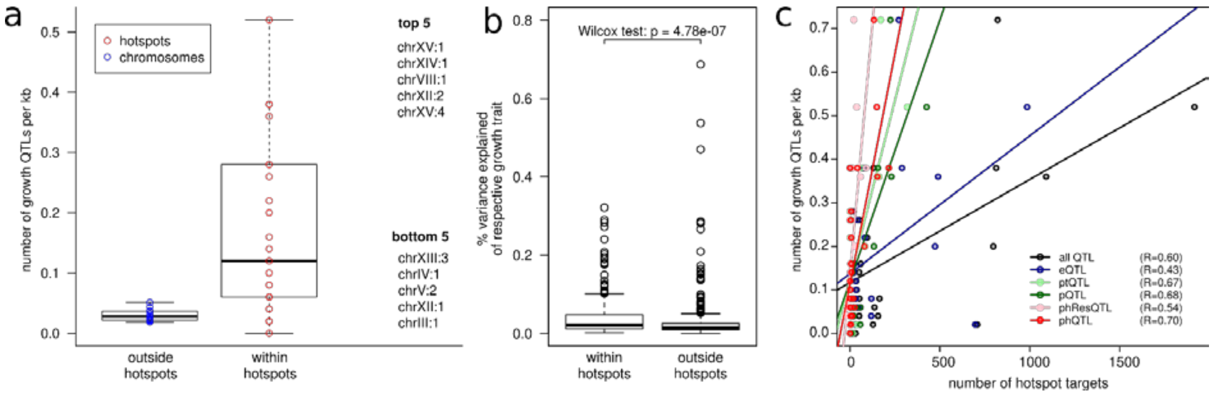


**Supplementary Figure S11** QTL hotspots for molecular traits also affect growth traits. Growth QTLs from Bloom *et al*. whose peaks were within 50 kb of the loci with the most targets in our hotspots were considered the same QTL. (**a**) Density of growth QTLs close to hotspots (“within hotspots”) compared to the rest of the genome (“outside hotspots”). Here, density refers to the number of growth QTLs (any growth condition) per kb. For hotspots, the circles represent individual hotspots (red), for the rest of the genome the circles represent individual chromosomes (blue). The hotspots with the five highest and lowest growth QTL density are listed on the right. (**b**) Distribution of the proportion of growth phenotype variance that was explained by growth QTLs that were close to hotspots, or not, respectively. (**c**) Relationship between the number of targets at each layer and the number of growth QTLs that were affected by each hotspot. Linear regressions and Pearson correlation coefficients are indicated. All correlations were significant with a corrected p-value below 0.05.

### Supplementary Tables

**Supplementary Table S1** Amount of QTL. Shown are the absolute number of associations between a QTL and a trait for each molecular layer and the number of traits that are affected by at least one QTL at FDR<10%.

| layer | Number of QTL | Affected traits |
|---|---|---|
| eQTL | 5776 | 4202 |
| ptQTL | 1327 | 1081 |
| pQTL | 2078 | 1438 |
| phResQTL | 466 | 389 |
| phQTL | 1595 | 1266 |

**Supplementary Table S2** Overview of QTL hotspots. Each region that was called as a QTL hotspot is shown as a row. For each hotspot the location, the number of targets on each layer and the significantly affected layers are given. If a causal gene was identified, the gene and the reference are shown.

| name | chr | Start (bp) | End (bp) | Start (bin) | End (Bin) | eQtl argets | ptQtl Targets | pQtl Targets | phResQtl Targets | phQtl Targets | eQtl Hotspot | ptQtl Hotspot | pQtl Hotspot | phResQtl Hotspot | phQtl Hotspot | Causal Gene | reference |
|---|---|---|---|---|---|---|---|---|---|---|---|---|---|---|---|---|---|
| chrI:1 | chrI | 1 | 40001 | 1 | 1 | 42 | 3 | 6 | 0 | 3 | TRUE | FALSE | FALSE | FALSE | FALSE | unknown | - |
| chrII:1 | chrII | 320001 | 360001 | 14 | 14 | 22 | 25 | 57 | 7 | 25 | FALSE | TRUE | TRUE | TRUE | TRUE | unknown | - |
| chrII:2 | chrII | 400001 | 640001 | 16 | 21 | 473 | 83 | 133 | 27 | 81 | TRUE | TRUE | TRUE | TRUE | TRUE | *AMN1* | PMID: 12897782 |
| chrIII:1 | chrIII | 80001 | 120001 | 28 | 28 | 19 | 10 | 17 | 1 | 4 | FALSE | FALSE | TRUE | FALSE | FALSE | *LEU2* | PMID: 11923494 |
| chrIV:1 | chrIV | 1280001 | 1320001 | 65 | 65 | 1 | 0 | 1 | 0 | 3 | FALSE | FALSE | FALSE | FALSE | TRUE | unknown | - |
| chrV:1 | chrV | 360001 | 440001 | 80 | 81 | 119 | 6 | 16 | 5 | 18 | TRUE | FALSE | FALSE | TRUE | TRUE | unknown | - |
| chrV:2 | chrV | 480001 | 576874 | 83 | 84 | 696 | 7 | 0 | 1 | 4 | TRUE | FALSE | FALSE | FALSE | FALSE | unknown | - |
| chrVII:1 | chrVII | 120001 | 160001 | 94 | 94 | 82 | 1 | 2 | 3 | 7 | TRUE | FALSE | FALSE | FALSE | FALSE | unknown | - |
| chrVII:2 | chrVII | 360001 | 400001 | 100 | 100 | 32 | 5 | 9 | 3 | 3 | FALSE | FALSE | FALSE | TRUE | FALSE | unknown | - |
| chrVIII:1 | chrVIII | 40001 | 120001 | 119 | 120 | 69 | 1 | 8 | 12 | 41 | TRUE | FALSE | FALSE | TRUE | TRUE | *GPA1* | PMID: 12897782 |
| chrVIII:2 | chrVIII | 240001 | 280001 | 124 | 124 | 33 | 1 | 1 | 0 | 2 | TRUE | FALSE | FALSE | FALSE | FALSE | unknown | - |
| chrX:1 | chrX | 280001 | 360001 | 149 | 150 | 118 | 6 | 20 | 1 | 11 | TRUE | FALSE | TRUE | FALSE | FALSE | unknown | - |
| chrXII:1 | chrXII | 480001 | 520001 | 188 | 188 | 31 | 35 | 57 | 0 | 5 | FALSE | TRUE | TRUE | FALSE | FALSE | unknown | - |
| chrXII:2 | chrXII | 560001 | 720001 | 190 | 193 | 289 | 64 | 155 | 89 | 216 | TRUE | TRUE | TRUE | TRUE | TRUE | *HAP1* | PMID: 11923494 |
| chrXIII:1 | chrXIII | 1 | 200001 | 202 | 206 | 490 | 161 | 230 | 58 | 152 | TRUE | TRUE | TRUE | TRUE | TRUE | unknown | - |
| chrXIII:2 | chrXIII | 240001 | 280001 | 208 | 208 | 27 | 9 | 16 | 3 | 5 | FALSE | TRUE | TRUE | FALSE | FALSE | unknown | - |
| chrXIII:3 | chrXIII | 520001 | 560001 | 215 | 215 | 8 | 15 | 8 | 0 | 0 | FALSE | TRUE | FALSE | FALSE | FALSE | unknown | - |
| chrXIII:4 | chrXIII | 880001 | 924431 | 224 | 224 | 31 | 0 | 2 | 0 | 1 | TRUE | FALSE | FALSE | FALSE | FALSE | unknown | - |
| chrXIV:1 | chrXIV | 400001 | 560001 | 235 | 238 | 985 | 319 | 426 | 37 | 149 | TRUE | TRUE | TRUE | TRUE | TRUE | *MKT1* | PMID: 11923494 |
| chrXV:1 | chrXV | 40001 | 200001 | 245 | 248 | 271 | 171 | 225 | 21 | 133 | TRUE | TRUE | TRUE | TRUE | TRUE | *IRA2* | PMID: 12897782 |
| chrXV:2 | chrXV | 240001 | 280001 | 250 | 250 | 19 | 3 | 16 | 0 | 1 | FALSE | FALSE | TRUE | FALSE | FALSE | unknown | - |
| chrXV:3 | chrXV | 440001 | 480001 | 255 | 255 | 46 | 3 | 4 | 0 | 5 | TRUE | FALSE | FALSE | FALSE | FALSE | unknown | - |
| chrXV:4 | chrXV | 520001 | 560001 | 257 | 257 | 70 | 3 | 4 | 0 | 0 | TRUE | FALSE | FALSE | FALSE | FALSE | unknown | - |
| chrXV:5 | chrXV | 600001 | 640001 | 259 | 259 | 22 | 5 | 14 | 2 | 8 | FALSE | FALSE | TRUE | FALSE | FALSE | unknown | - |
| chrXVI:1 | chrXVI | 440001 | 480001 | 282 | 282 | 9 | 1 | 2 | 2 | 6 | FALSE | FALSE | FALSE | TRUE | FALSE | *DIG1* | PMID: 12897782 |

**Supplementary Table S3** GO-enrichments of genes targeted by eQTL with similar effects on transcript and protein levels (‘similar’ in Figure 3b).

| ontology | GO.ID | Term | Annotated | Significant | Expected | p-value |
|---|---|---|---|---|---|---|
| BP | GO:0030150 | protein import into mitochondrial matrix | 15 | 14 | 8.76 | 0.0036 |
| BP | GO:0097502 | Mannosylation | 21 | 16 | 12.27 | 0.0059 |
| BP | GO:0031505 | fungal-type cell wall organization | 61 | 46 | 35.64 | 0.0064 |
| BP | GO:0006696 | ergosterol biosynthetic process | 21 | 18 | 12.27 | 0.0073 |
| BP | GO:0016485 | protein processing | 17 | 16 | 9.93 | 0.0091 |
| MF | GO:0005525 | GTP binding | 45 | 36 | 26.29 | 0.0017 |
| MF | GO:0016887 | ATPase activity | 110 | 63 | 64.27 | 0.0099 |

**Supplementary Table S4** GO-enrichments of genes targeted by eQTL with strong effects on transcript levels and weak effects on protein levels ('buffered' in Figure 3b).

| ontology | GO.ID | Term | Annotated | Significant | Expected | p-value |
|---|---|---|---|---|---|---|
| BP | GO:0000463 | maturation of LSU-rRNA from tricistronic... | 23 | 17 | 4.69 | 3.6E-09 |
| BP | GO:0006364 | rRNA processing | 134 | 66 | 27.35 | 0.000004 |
| BP | GO:0002181 | cytoplasmic translation | 75 | 29 | 15.31 | 0.000039 |
| BP | GO:0000447 | endonucleolytic cleavage in ITS1 to sepa... | 20 | 12 | 4.08 | 0.00011 |
| BP | GO:0000466 | maturation of 5.8S rRNA from tricistroni... | 41 | 21 | 8.37 | 0.00036 |
| BP | GO:0000472 | endonucleolytic cleavage to generate mat... | 18 | 10 | 3.67 | 0.00101 |
| BP | GO:0000027 | ribosomal large subunit assembly | 21 | 11 | 4.29 | 0.00108 |
| BP | GO:0042797 | tRNA transcription from RNA polymerase I... | 13 | 8 | 2.65 | 0.00137 |
| BP | GO:0000480 | endonucleolytic cleavage in 5'-ETS of tr... | 16 | 9 | 3.27 | 0.00163 |
| BP | GO:0006259 | DNA metabolic process | 126 | 20 | 25.72 | 0.00189 |
| BP | GO:0006526 | arginine biosynthetic process | 9 | 6 | 1.84 | 0.00334 |
| BP | GO:0006360 | transcription from RNA polymerase I prom... | 37 | 9 | 7.55 | 0.00517 |
| BP | GO:0000055 | ribosomal large subunit export from nucl... | 16 | 8 | 3.27 | 0.0077 |
| BP | GO:0042254 | ribosome biogenesis | 196 | 88 | 40 | 0.00774 |
| BP | GO:0031668 | cellular response to extracellular stimu... | 32 | 12 | 6.53 | 0.00824 |
| MF | GO:0001056 | RNA polymerase III activity | 11 | 8 | 2.25 | 0.00026 |
| MF | GO:0003723 | RNA binding | 275 | 82 | 56.13 | 0.00048 |
| MF | GO:0001054 | RNA polymerase I activity | 11 | 7 | 2.25 | 0.00216 |
| MF | GO:0042134 | rRNA primary transcript binding | 5 | 4 | 1.02 | 0.00718 |
| MF | GO:0019200 | carbohydrate kinase activity | 7 | 5 | 1.43 | 0.00837 |
| CC | GO:0005730 | Nucleolus | 144 | 78 | 29.39 | 2.7E-18 |
| CC | GO:0030687 | preribosome, large subunit precursor | 47 | 30 | 9.59 | 6.5E-11 |
| CC | GO:0022625 | cytosolic large ribosomal subunit | 36 | 18 | 7.35 | 0.000061 |
| CC | GO:0005666 | DNA-directed RNA polymerase III complex | 11 | 8 | 2.25 | 0.00026 |
| CC | GO:0005654 | nucleoplasm | 96 | 27 | 19.59 | 0.00054 |
| CC | GO:0032040 | small-subunit processome | 27 | 13 | 5.51 | 0.00106 |
| CC | GO:0005736 | DNA-directed RNA polymerase I complex | 11 | 7 | 2.25 | 0.00216 |
| CC | GO:0070545 | PeBoW complex | 3 | 3 | 0.61 | 0.00845 |

**Supplementary Table S5** GO-enrichments of genes targeted by eQTL with stronger effects on protein levels than on transcript levels (‘enhanced’ in Figure 3b).

| ontology | GO.ID | Term | Annotated | Significant | Expected | p-value |
|---|---|---|---|---|---|---|
| BP | GO:0032543 | mitochondrial translation | 76 | 37 | 6.71 | 1.9E-20 |
| BP | GO:0019419 | sulfate reduction | 3 | 3 | 0.26 | 0.00068 |
| BP | GO:0019344 | cysteine biosynthetic process | 8 | 4 | 0.71 | 0.0031 |
| BP | GO:0006839 | mitochondrial transport | 56 | 7 | 4.95 | 0.00548 |
| BP | GO:0070814 | hydrogen sulfide biosynthetic process | 5 | 3 | 0.44 | 0.00592 |
| BP | GO:0000947 | amino acid catabolic process to alcohol ... | 5 | 3 | 0.44 | 0.00592 |
| BP | GO:0005975 | carbohydrate metabolic process | 127 | 10 | 11.22 | 0.006 |
| MF | GO:0003735 | structural constituent of ribosome | 102 | 34 | 9.01 | 4.6E-13 |
| MF | GO:0016744 | transferase activity, transferring aldeh... | 4 | 3 | 0.35 | 0.0025 |
| MF | GO:0004022 | alcohol dehydrogenase (NAD) activity | 4 | 3 | 0.35 | 0.0025 |
| CC | GO:0005762 | mitochondrial large ribosomal subunit | 32 | 20 | 2.83 | 2.5E-14 |
| CC | GO:0005763 | mitochondrial small ribosomal subunit | 26 | 14 | 2.3 | 3.9E-09 |
| CC | GO:0005737 | cytoplasm | 1615 | 160 | 142.63 | 0.0006 |
| CC | GO:0005759 | mitochondrial matrix | 137 | 46 | 12.1 | 0.0011 |
| CC | GO:0005739 | mitochondrion | 531 | 93 | 46.89 | 0.0048 |

**Supplementary Table S6** GO-enrichments of genes targeted by ptQTL (‘protein only’ in Figure 3b).

| ontology | GO.ID | Term | Annotated | Significant | Expected | p-value |
|---|---|---|---|---|---|---|
| BP | GO:0002181 | cytoplasmic translation | 75 | 52 | 37.2 | 0.000047 |
| BP | GO:0055114 | oxidation-reduction process | 239 | 145 | 118.53 | 0.0076 |
| BP | GO:0006099 | tricarboxylic acid cycle | 19 | 15 | 9.42 | 0.0085 |
| CC | GO:0022625 | cytosolic large ribosomal subunit | 36 | 27 | 17.85 | 0.0015 |
| CC | GO:0042645 | mitochondrial nucleoid | 16 | 13 | 7.94 | 0.0095 |

**Supplementary Table S7** QTL affecting growth under various conditions were taken from Ref. 20. For growth QTL, we used the reported peak QTL positions. For our hotspots, we took the midpoint of the peak QTLs of each layer (i.e., the loci that regulated most traits on each of the molecular layers). If growth QTL and hotspot peak were within 50 kb of each other, we considered them to be the same QTL. Listed below are the growth QTL affected by hotspots that were mentioned in the text. Hotspots that affected only expression typically affected fewer growth traits than hotspots that affected many phospho-traits.

| chrV:2 | chrVIII:1 | chrXII:1 | chrXII:2 | chrXIV:1 | chrXV:1 |
|---|---|---|---|---|---|
| Menadione | 5-Fluorocytosine | Cobalt_Chloride | 4-Hydroxybenzaldehyde | 4-Hydroxybenzaldehyde | 4NQO |
| | 5-Fluorouracil | | 4NQO | 4NQO | 5-Fluorocytosine |
| | Caffeine | | Cadmium_Chloride | 5-Fluorocytosine | 5-Fluorouracil |
| | Congo_red | | Calcium_Chloride | 5-Fluorouracil | 6-Azauracil |
| | Cycloheximide | | Cisplatin | Caffeine | Calcium_Chloride |
| | Diamide | | Cobalt_Chloride | Calcium_Chloride | Cisplatin |
| | E6_Berbamine | | Copper | Cisplatin | Cobalt_Chloride |
| | Formamide | | Cycloheximide | Copper | Congo_red |
| | Indoleacetic_Acid | | E6_Berbamine | Cycloheximide | Copper |
| | Menadione | | Ethanol | E6_Berbamine | Cycloheximide |
| | SDS | | Formamide | Ethanol | E6_Berbamine |
| | YNB | | Lithium_Chloride | Formamide | Ethanol |
| | YNB:ph3 | | Manganese_Sulfate | Hydrogen_Peroxide | Formamide |
| | YNB:ph8 | | Mannose | Indoleacetic_Acid | Galactose |
| | YPD | | Neomycin | Magnesium_Chloride | Hydrogen_Peroxide |
| | YPD:15C | | Tunicamycin | Raffinose | Lactate |
| | Zeocin | | YNB | Trehalose | Lactose |
| | | | YPD | Tunicamycin | Lithium_Chloride |
| | | | Zeocin | Xylose | Magnesium_Chloride |
| | | | | YNB | Maltose |
| | | | | YNB:ph3 | Manganese_Sulfate |
| | | | | YPD | Mannose |
| | | | | YPD:37C | Neomycin |
| | | | | YPD:4C | Paraquat |
| | | | | | Raffinose |
| | | | | | SDS |
| | | | | | Sorbitol |
| | | | | | Trehalose |
| | | | | | Xylose |
| | | | | | YNB |
| | | | | | YNB:ph3 |
| | | | | | YPD |
| | | | | | YPD:15C |
| | | | | | YPD:37C |
| | | | | | YPD:4C |
| | | | | | Zeocin |

**Supplementary Table S8** Connection of phQTL- and/or phResQTL-targets of chrXV:1 to cAMP-signaling. As protein levels are not available for all genes with available phosphopeptide levels, we coded non available levels as NA with respect to pQTL and phResQTL.

| systematic name | standard name | connection to cAMP signaling | reference | has phResQTL | has phQTL | has pQTL |
|---|---|---|---|---|---|---|
| YAL034C | FUN19 | NA | NA | NA | 1 | NA |
| YBL007C | SLA1 | Phosphorylation is dependent on PKA activity | PMID: 21177495 | 1 | 1 | 0 |
| YBL024W | NCL1 | NA | NA | NA | 1 | NA |
| YBL032W | HEK2 | NA | NA | 1 | 1 | 0 |
| YBL054W | TOD6 | NA | NA | NA | 1 | NA |
| YBL061C | SKT5 | Phosphorylation is dependent on PKA activity | PMID: 21177495 | NA | 1 | NA |
| YBL085W | BOI1 | BOI1 physically interacts with CYR1 | PMID: 16429126 | NA | 1 | NA |
| YBR023C | CHS3 | NA | NA | NA | 1 | NA |
| YBR114W | RAD16 | NA | NA | NA | 1 | NA |
| YBR214W | SDS24 | Phosphorylation is dependent on PKA activity | PMID: 21177495 | 0 | 1 | 1 |
| YCL014W | BUD3 | Target of Tpk1 | PMID: 16319894 | NA | 1 | NA |
| YCR091W | KIN82 | NA | NA | NA | 1 | NA |
| YDL019C | OSH2 | NA | NA | NA | 1 | NA |
| YDL054C | MCH1 | NA | NA | NA | 1 | NA |
| YDL113C | ATG20 | Member of the CVT-pathway whose activity is partially regulated through PKA-signaling | PMID: 16172400 | NA | 1 | NA |
| YDL146W | LDB17 | NA | NA | NA | 1 | NA |
| YDL222C | FMP45 | Involved in eisosome assembly with proteins whose phosphorylation is dependent on PKA-activity (LSP1, EIS1, YGR130C, SEG2) | PMID: 21177495, PMID: 11784867 | 0 | 1 | 1 |
| YDL223C | HBT1 | Phosphorylation is dependent on PKA activity | PMID: 21177495 | 1 | 1 | 1 |
| YDR006C | SOK1 | Target of Yak1, overexpression can rescue Tpk1-3 deletion | PMID: 8065298 | NA | 1 | NA |
| YDR074W | TPS2 | Forms a complex with other targets of PKA-signaling (TPS3, TSL1) | PMID: 21177495, PMID: 8879249 | 0 | 1 | 1 |
| YDR135C | YCF1 | Phosphorylation is dependent on PKA activity | PMID: 21177495 | 1 | 1 | 0 |
| YDR207C | UME6 | Phosphorylation is dependent on PKA activity , target of Rim11 | PMID: 16319894 | NA | 1 | NA |
| YDR216W | ADR1 | Phosphorylation is dependent on PKA activity | PMID: 2644045 | 0 | 1 | 1 |
| YDR293C | SSD1 | Phosphorylation is dependent on PKA activity | PMID: 21177495 | NA | 1 | NA |
| YDR310C | SUM1 | NA | NA | 1 | 1 | 0 |
| YDR326C | YSP2 | NA | NA | NA | 1 | NA |
| YDR409W | SIZ1 | NA | NA | NA | 1 | NA |
| YDR508C | GNP1 | NA | NA | NA | 1 | NA |
| YEL015W | EDC3 | Phosphorylation is dependent on PKA activity | PMID: 16319894 | 0 | 1 | 0 |
| YER054C | GIP2 | Phosphorylation is dependent on PKA activity | PMID: 21177495 | NA | 1 | NA |
| YER129W | SAK1 | NA | NA | NA | 1 | NA |
| YFL014W | HSP12 | Phosphorylation is dependent on PKA activity | PMID: 21177495 | 0 | 1 | 1 |
| YFR017C | IGD1 | Phosphorylation is dependent on PKA activity | PMID: 21177495 | NA | 1 | NA |
| YGL201C | MCM6 | Forms a complex with MCM4, a target of PKA | PMID: 16319894, PMID: 16824194 | NA | 1 | NA |
| YGR054W | YGR054W | NA | NA | NA | 1 | NA |
| YGR086C | PIL1 | Involved in eisosome assembly with proteins whose phosphorylation is | PMID: 21177495, PMID: 25057013 | 0 | 1 | 1 |

| systematic name | standard name | connection to cAMP signaling | reference | has phResQTL | has phQTL | has pQTL |
|---|---|---|---|---|---|---|
| | | dependent on PKA-activity (LSP1, EIS1, YGR130C, SEG2) | | | | |
| YGR097W | ASK10 | NA | NA | NA | 1 | NA |
| YGR116W | SPT6 | Associates with TPK2 | PMID: 20489023 | 1 | 0 | 0 |
| YGR125W | YGR125W | Phosphorylation is dependent on PKA activity | PMID: 21177495 | NA | 1 | NA |
| YGR130C | YGR130C | Phosphorylation is dependent on PKA activity | PMID: 21177495 | 0 | 1 | 1 |
| YGR145W | ENP2 | NA | NA | NA | 1 | NA |
| YGR159C | NSR1 | Phosphorylation is dependent on PKA activity | PMID: 16319894 | 1 | 1 | 0 |
| YGR185C | TYS1 | NA | NA | NA | 1 | NA |
| YGR261C | APL6 | NA | NA | NA | 1 | NA |
| YGR270W | YTA7 | NA | NA | NA | 1 | NA |
| YHL007C | STE20 | NA | NA | 1 | 1 | 0 |
| YHR064C | SSZ1 | NA | NA | 0 | 1 | 0 |
| YHR097C | YHR097C | Phosphorylation is dependent on PKA activity | PMID: 21177495 | 0 | 1 | 1 |
| YHR103W | SBE22 | NA | NA | NA | 1 | NA |
| YIL105C | SLM1 | Involved in eisosome assembly with proteins whose phosphorylation is dependent on PKA-activity (LSP1, EIS1, YGR130C, SEG2) | PMID: 21177495, PMID: 19064668 | 0 | 1 | 1 |
| YIL122W | POG1 | Phosphorylation is dependent on PKA activity | PMID: 21177495 | NA | 1 | NA |
| YIL135C | VHS2 | Phosphorylation is dependent on PKA activity | PMID: 21177495 | 0 | 1 | 0 |
| YIL136W | OM45 | NA | NA | 0 | 1 | 1 |
| YIR038C | GTT1 | NA | NA | 0 | 1 | 1 |
| YJL042W | MHP1 | NA | NA | 0 | 1 | 0 |
| YJL123C | MTC1 | NA | NA | 1 | 1 | 0 |
| YJL141C | YAK1 | Yak1 is phosphorylates Hsf1 and Msn2/4 downstream of the PKA-complex | PMID: 18793336 | 0 | 1 | 1 |
| YJR001W | AVT1 | Phosphorylation is dependent on PKA activity | PMID: 16172400, PMID: 16319894 | 0 | 1 | 0 |
| YJR092W | BUD4 | Physically interacts with PKA-target BUD3 | PMID: 2048902 | NA | 1 | NA |
| YKL035W | UGP1 | Phosphorylation is dependent on PKA activity | PMID: 21177495 | 0 | 1 | 1 |
| YKL062W | MSN4 | The localization of Msn4 is regulated through the PKA-complex and possibly Yak1. | PMID: 9472026, PMID: 18793336 | 0 | 1 | 1 |
| YKL105C | SEG2 | Phosphorylation is dependent on PKA activity | PMID: 21177495 | NA | 1 | NA |
| YKL112W | ABF1 | NA | NA | NA | 1 | NA |
| YKL160W | ELF1 | Phosphorylation is dependent on PKA activity | PMID: 16319894 | 0 | 1 | 0 |
| YKL175W | ZRT3 | NA | NA | NA | 1 | NA |
| YKR019C | IRS4 | NA | NA | NA | 1 | NA |
| YKR093W | PTR2 | NA | NA | NA | 1 | NA |
| YLL013C | PUF3 | NA | NA | NA | 1 | NA |
| YLL028W | TPO1 | Localization of TPO1 is affected by phosphorylation through PKA | PMID: 15637075 | NA | 1 | NA |
| YLL061W | MMP1 | NA | NA | NA | 1 | NA |
| YLR006C | SSK1 | NA | NA | NA | 1 | NA |
| YLR044C | PDC1 | Regulated through phosphorylation in response to changes in glucose levels bin Sit4, physically interacts also with Bcy1, Target of TPK3 | PMID: 23692511, PMID: 25065647, PMID: 16319894 | NA | 1 | NA |
| YLR096W | KIN2 | NA | NA | NA | 1 | NA |
| YLR187W | SKG3 | NA | NA | NA | 1 | NA |

| systematic name | standard name | connection to cAMP signaling | reference | has phResQTL | has phQTL | has pQTL |
|---|---|---|---|---|---|---|
| YLR206W | ENT2 | NA | NA | NA | 1 | NA |
| YLR219W | MSC3 | Involved in eisosome assembly with proteins whose phosphorylation is dependent on PKA-activity (LSP1, EIS1, YGR130C, SEG2) | PMID: 21177495, PMID: 19037108 | 1 | 1 | 0 |
| YLR249W | YEF3 | NA | NA | 1 | 0 | 0 |
| YLR257W | YLR257W | Phosphorylation is dependent on PKA activity | PMID: 21177495 | 1 | 1 | 0 |
| YLR310C | CDC25 | CDC25 promotes the GTP-bound form of Ras | PMID: 3545497, PMID: 2188363 | NA | 1 | NA |
| YLR399C | BDF1 | Phosphorylation is dependent on PKA activity | PMID: 21177495 | NA | 1 | NA |
| YLR429W | CRN1 | Phosphorylation is dependent on PKA activity | PMID: 21177495 | NA | 1 | NA |
| YML061C | PIF1 | NA | NA | NA | 1 | NA |
| YML070W | DAK1 | NA | NA | 0 | 1 | 1 |
| YML100W | TSL1 | Phosphorylation is dependent on PKA activity | PMID: 21177495 | 0 | 1 | 1 |
| YML127W | RSC9 | NA | NA | 1 | 1 | 0 |
| YMR011W | HXT2 | NA | NA | 1 | 1 | 0 |
| YMR031C | EIS1 | Phosphorylation is dependent on PKA activity | PMID: 21177495 | 1 | 1 | 1 |
| YMR105C | PGM2 | Phosphorylation is dependent on PKA activity | PMID: 21177495 | 0 | 1 | 1 |
| YMR131C | RRB1 | NA | NA | NA | 1 | NA |
| YMR139W | RIM11 | Involved in downstream signaling of PKA | PMID: 15282298 | 0 | 1 | 1 |
| YMR196W | YMR196W | Phosphorylation is dependent on PKA activity | PMID: 16319894 | 0 | 1 | 1 |
| YMR261C | TPS3 | Phosphorylation is dependent on PKA activity | PMID: 21177495 | 0 | 1 | 0 |
| YNL074C | MLF3 | Phosphorylation is dependent on PKA activity | PMID: 16319894 | 0 | 1 | 0 |
| YNL098C | RAS2 | GTP-bound Ras promotes cAMP-production by CYR1 | PMID: 3891097 | 0 | 1 | 0 |
| YNL101W | AVT4 | Phosphorylation is dependent on PKA activity | PMID: 16319894 | NA | 1 | NA |
| YNL106C | INP52 | NA | NA | NA | 1 | NA |
| YNL113W | RPC19 | NA | NA | 0 | 1 | 0 |
| YNL173C | MDG1 | Phosphorylation is dependent on PKA activity | PMID: 21177495 | 1 | 1 | 0 |
| YNL175C | NOP13 | NA | NA | 1 | 1 | 0 |
| YNL183C | NPR1 | NPR1 deletions change RAS-signaling activity | PMID: 11238915 | NA | 1 | NA |
| YNL229C | URE2 | NA | NA | NA | 1 | NA |
| YNL230C | ELA1 | NA | NA | NA | 1 | NA |
| YNL234W | YNL234W | NA | NA | NA | 1 | NA |
| YNL267W | PIK1 | NA | NA | NA | 1 | NA |
| YNL268W | LYP1 | NA | NA | NA | 1 | NA |
| YNL274C | GOR1 | NA | NA | 0 | 1 | 1 |
| YNL278W | CAF120 | NA | NA | NA | 1 | NA |
| YNL298W | CLA4 | NA | NA | 1 | 0 | 0 |
| YNL304W | YPT11 | NA | NA | NA | 1 | NA |
| YNR014W | YNR014W | Phosphorylation is dependent on PKA activity | PMID: 21177495 | NA | 1 | NA |
| YNR019W | ARE2 | Phosphorylation is dependent on PKA activity | PMID: 16319894 | 0 | 1 | 0 |
| YOL004W | SIN3 | Member of the RPD3L complex of which also Ume6, a target of PKA, is a part of | PMID: 16319894 | NA | 1 | NA |
| YOL078W | AVO1 | NA | NA | NA | 1 | NA |
| YOL081W | IRA2 | Phosphorylation is dependent on PKA activity | PMID: 21177495 | NA | 1 | NA |

| systematic name | standard name | connection to cAMP signaling | reference | has phResQTL | has phQTL | has pQTL |
|---|---|---|---|---|---|---|
| YOL082W | ATG19 | Phosphorylation is dependent on PKA activity | PMID: 21177495 | NA | 1 | NA |
| YOL109W | ZEO1 | NA | NA | 1 | 1 | 1 |
| YOL130W | ALR1 | Phosphorylation is dependent on PKA activity | PMID: 16319894 | NA | 1 | NA |
| YOL137W | BSC6 | NA | NA | NA | 1 | NA |
| YOR028C | CIN5 | NA | NA | NA | 1 | NA |
| YOR066W | MSA1 | NA | NA | NA | 1 | NA |
| YOR161C | PNS1 | NA | NA | NA | 1 | NA |
| YOR227W | HER1 | NA | NA | NA | 1 | NA |
| YOR353C | SOG2 | NA | NA | NA | 1 | NA |
| YOR371C | GPB1 | promotes proteolysis of Ira2 | PMID: 20160012 | NA | 1 | NA |
| YPL004C | LSP1 | Phosphorylation is dependent on PKA activity | PMID: 21177495 | 0 | 1 | 1 |
| YPL019C | VTC3 | Phosphorylation is dependent on PKA activity | PMID: 21177495 | 0 | 1 | 1 |
| YPL085W | SEC16 | Phosphorylation is dependent on PKA activity | PMID: 21177495 | 1 | 1 | 0 |
| YPL115C | BEM3 | NA | NA | NA | 1 | NA |
| YPL186C | UIP4 | NA | NA | NA | 1 | NA |
| YPL247C | YPL247C | Colocalizes with Tpk2 and Tpk3 | PMID: 2048902 | 0 | 1 | 1 |
| YPR016C | TIF6 | NA | NA | 0 | 1 | 0 |
| YPR019W | MCM4 | Phosphorylation is dependent on PKA activity | PMID: 16319894 | NA | 1 | NA |
| YPR036W-A | SPO24 | NA | NA | 0 | 1 | 0 |
| YPR117W | YPR117W | NA | NA | NA | 1 | NA |
| YPR149W | NCE102 | Involved in eisosome assembly with proteins whose phosphorylation is dependent on PKA-activity (LSP1, EIS1, YGR130C, SEG2) | PMID: 21177495, PMID: 19064668 | 0 | 1 | 1 |
| YPR161C | SGV1 | Phosphorylation is dependent on PKA activity | PMID: 21177495 | 1 | 1 | 0 |

**Supplementary Table S9** Targets of the Gpa1/Ste20 hotspot on ChrVIII:1 which could be functionally linked to Ste20 and/or Gpa1 through literature research. Related to Figure 5.

| systematic name | standard name | eQTL FC | ptQTL FC | pQTL FC | phResQTL FC | phQTL FC | local QTL | Ste20 link | references |
|---|---|---|---|---|---|---|---|---|---|
| YAL031C | GIP4 | 0.09 | NA | NA | NA | NA | 0 | Interacts with Glc7, which, together with Reg1, acts as a GTPase for Gpa1. | PMID: 16537909 |
| YCL027W | FUS1 | 0.89 | NA | NA | NA | NA | 0 | FUS1 is transcribed upon activation by Ste12, which is a transcription factor that is activated via the pheromone MAP kinase cascade dependent of Ste20. | PMID: 17604854, PMID: 1903837 |
| YCL055W | KAR4 | 0.40 | NA | NA | NA | NA | 0 | Transcriptional target of the transcription factor Ste12, which is activated though MAP kinase cascade. | PMID: 17604854, PMID: 16522208 |
| YDL240W | LRG1 | 0.09 | NA | NA | NA | NA | 0 | Acts as a GTPase-activating protein (GAP) for Cdc42, inhibiting it. | PMID: 27704052, PMID: 11591390 |
| YDR379W | RGA2 | 0.07 | NA | NA | NA | NA | 0 | Rga2 is a GTPase associated with Cdc42, upstream of Ste20. | PMID: 12455995 |
| YDR461W | MFA1 | 0.58 | NA | NA | NA | NA | 0 | Transcriptionally regulated by Ste12 which is a transcription factor driving the pheromone response and that is activated by the MAPK pathway. | PMID: 2659433, PMID: 17604854 |
| YDR530C | APA2 | 0.10 | NA | NA | NA | NA | 0 | Transcriptionally regulated by Ste12 which is a transcription factor driving the pheromone response and that is activated by the MAPK pathway. | PMID: 16522208, PMID: 12732146 |
| YFL047W | RGD2 | 0.16 | NA | NA | NA | NA | 0 | Acts as a GAP for Cdc42, repressing the activation of Ste20. | PMID: 24062589, PMID: 11591390 |
| YGR189C | CRH1 | -0.07 | NA | NA | NA | NA | 0 | Is activated by the MAPK pathway via Hog1 and the cell wall integrity pathway. | PMID: 16522208, PMID: 19234305, PMID: 18184748 |
| YHR005C | GPA1 | 0.19 | NA | NA | NA | NA | 1 | Gpa1 is associated with Ste4 and Ste18, and is dissociated upon GDP to GTP exchange when pheromone signal is received. This triggers the activation of Ste20. Gpa1 also interacts with Fus3 to antagonize its action. | PMID: 2536595, PMID: 9832519, PMID: 10712512, PMID: 12556475 |
| YIL117C | PRM5 | 0.56 | NA | NA | NA | NA | 0 | Regulated by Ste12 and known to be pheromone-regulated. | PMID: 10535956, PMID: 11062271 |
| YJL085W | EXO70 | 0.05 | NA | NA | NA | NA | 0 | Exo70 is a direct effector for Cdc42, which is just upstream of Ste20 in the MAPK pathway. | PMID: 19955214 |
| YJL157C | FAR1 | 0.30 | NA | NA | NA | 0.75, 0.78 | 0 | Interacts directly with Gpa1. Phosphorylated by Fus3 which is a downstream kinase of Ste20 in the pheromone response pathway. | PMID: 12029138, PMID: 18261907, PMID: 8500168, PMID: 15690603 |
| YJR086W | STE18 | 0.12 | NA | NA | NA | NA | 0 | Ste4 and Ste18 form a dimer that gets released by Gpa1 upon GDP to GTP exchange when a pheromone signal is received. | PMID: 2536595, PMID: 7834739 |

| systematic name | standard name | eQTL FC | ptQTL FC | pQTL FC | phResQTL FC | phQTL FC | local QTL | Ste20 link | references |
|---|---|---|---|---|---|---|---|---|---|
| YKL189W | HYM1 | 0.17 | NA | NA | NA | NA | 0 | Transcriptionally regulated by Ste12 which is a transcription factor driving the pheromone response and that is activated by the Ste20 MAPK pathway. | PMID: 16522208 |
| YLR452C | SST2 | 0.54 | NA | NA | NA | NA | 0 | GTPase of Gpa1, which is a key effector of the pheromone response pathway, upstream of Ste20. | PMID: 9537998 |
| YML046W | PRP39 | 0.12 | NA | NA | NA | NA | 0 | Transcriptionally regulated by Ste12p which is a transcription factor driving the pheromone response and that is activated by the MAPK pathway. | PMID: 16522208 |
| YMR065W | KAR5 | 0.27 | NA | NA | NA | NA | 0 | Regulated by Hog1, the exact mechanism remains unknown. | PMID: 12052881, PMID: 16522208 |
| YMR232W | FUS2 | 0.46 | NA | NA | NA | NA | 0 | Transcriptionally regulated by Ste12 which is a transcription factor driving the pheromone response and that is activated by the MAPK pathway. | PMID: 16522208, PMID: 11166190 |
| YNL279W | PRM1 | 0.79 | NA | NA | NA | NA | 0 | Transcriptionally regulated by Ste12 which is a transcription factor driving the pheromone response and that is activated by the MAPK pathway. | PMID: 11062271, PMID: 11166190, PMID: 16522208 |
| YNR044W | AGA1 | 0.75 | NA | NA | NA | NA | 0 | Transcriptionally regulated by Ste12 which is a transcription factor driving the pheromone response and that is activated by the MAPK pathway. | PMID: 11166190, PMID: 15690603, PMID: 2072914 |
| YOR054C | VHS3 | 0.13 | NA | NA | NA | NA | 0 | Transcriptionally controlled by the Hog1 pathway. Hog1 is the main effector of the osmotic shock response which is downstream of Ste20. | PMID: 25904326 |
| YOR219C | STE13 | 0.11 | NA | NA | NA | NA | 0 | Transcription is induced by a-factor, likely through Ste12. | PMID: 2685554 |
| YPL049C | DIG1 | 0.10 | NA | 0.18 | NA | 0.5, 0.77, 0.37 | 0 | Phosphorylated by Fus3 which relieves the inhibition of Ste12. Fus3 is a downstream MAP kinase of Ste20. | PMID: 17604854, PMID: 8918885, PMID: 9094309 |
| YPL156C | PRM4 | 0.30 | NA | NA | NA | NA | 0 | Pheromone-activated, likely regulated by Ste12. | PMID: 11062271, PMID: 11166190, PMID: 16522208 |
| YPR115W | RGC1 | 0.16 | NA | NA | NA | 0.16, 0.28 | 0 | Direct phosphorylation target of Hog1. Hog1 is the main effector of the osmotic shock response which is downstream of Ste20. | PMID: 24298058, PMID: 12142009, PMID: 10970855 |
| YBR083W | TEC1 | 0.20 | NA | NA | NA | NA | 0 | Downstream target of Ste20, Ste11, and Ste7. | PMID: 17118154 |
| YDR055W | PST1 | -0.08 | NA | NA | NA | NA | 0 | Transcriptional target of Tec1 and Ste12, which are downstream of Ste20 in the pheromone response pathway. | PMID: 10535956, PMID: 11062271 |
| YDR085C | AFR1 | 0.24 | NA | NA | NA | NA | 0 | Afr1 is regulated by Ste12 and Msn4, known to genetically interact with Ste2, and binds to Boi1, Boi2, and Glc7. All these proteins are downstream of Ste20. | PMID: 16522208, PMID: 11743162, PMID: 19841731, PMID: 18552279, |

| systematic name | standard name | eQTL FC | ptQTL FC | pQTL FC | phResQTL FC | phQTL FC | local QTL | Ste20 link | references |
|---|---|---|---|---|---|---|---|---|---|
| | | | | | | | | | PMID: 20489023, PMID: 20093466 |
| YGL116W | CDC20 | 0.10 | NA | NA | NA | NA | 0 | Target of Msn4, Cla4 and Ste20. | PMID: 21329885, PMID: 12642613 |
| YHL022C | SPO11 | 0.64 | NA | NA | NA | NA | 0 | Pheromone-induced (i.e., downstream of Gpa1/Ste20) and involved in recombination during meiosis. | PMID: 9456310 |
| YHL027W | RIM101 | 0.10 | NA | NA | NA | NA | 0 | Been shown to regulate the filamentous growth MAPK pathway (Kss1 pathway), which is regulated by Ste20. | PMID: 20333241 |
| YPR005C | HAL1 | 0.14 | NA | NA | NA | 0.45 | 0 | Transcriptional target of Ste12. Shown to have a stronger response to pheromone in Fus3 knockouts than in wild type. In addition, potentially downstream of the Hog1 pathway. | PMID: 16522208, PMID: 11525741, PMID: 12040128 |
| YBL016W | FUS3 | NA | NA | 0.30 | NA | NA | 0 | Fus3 is a component of the MAP kinase cascade activated downstream of Ste20. | PMID: 11525741 |
| YHR007C | ERG11 | NA | NA | -0.31 | NA | NA | 1 | Sterol metabolism is controlled by Ste20 via direct interaction with Erg11. Altogether this plays a role in cell polarity. | PMID: 17895367 |
| YIL016W | SNL1 | NA | -0.64 | -0.14 | NA | NA | 0 | Stimulated by pheromones, therefore likely related to Gpa1/Ste20 pathway. | PMID: 24121774 |
| YBL061C | SKT5 | NA | NA | NA | NA | 0.24 | 0 | Target of the transcription factor Ste12, which is downstream of Ste20. | PMID: 16522208, PMID: 12732146 |
| YBR059C | AKL1 | NA | NA | NA | NA | 0.15 | 0 | Interacts with Ste50 and Bem1, which are direct interactors of Ste20. | PMID: 19269370, PMID: 22875988 |
| YDR028C | REG1 | NA | NA | NA | NA | -0.15 | 0 | Cross talk between Cdc42/Ste20 and Reg1/Glc7. | PMID: 24003253 |
| YDR309C | GIC2 | NA | NA | NA | NA | 0.34, 0.19 | 0 | Cdc42 effector like Ste20, can promote mitotic exit independently of Ste20 via the same pathway. | PMID: 14734533, PMID: 9367979 |
| YDR480W | DIG2 | NA | NA | NA | NA | 0.35 | 0 | Part of the Ste12/Dig1/Dig2 transcription factor complex, which is one of the final transcription factors of the MAP kinase pathway for pheromone response. | PMID: 15690603, PMID: 17604854, PMID: 9841672 |
| YER114C | BOI2 | NA | NA | NA | NA | -0.2 | 0 | Forms a scaffold together with Bem1, for the binding of Ste20 and Cdc42. | PMID: 23785492 |
| YJR001W | AVT1 | NA | NA | NA | 0.62 | 0.28 | 0 | Regulated by Msn2, which is downstream from Hog1, which is downstream of Ste20. | PMID: 16522208 |
| YKL062W | MSN4 | NA | NA | NA | -1.07 | -0.44 | 0 | Downstream of the kinase Hog1 in the osmotic response pathway. Hog1 is the main effector of the osmotic shock response which is downstream of Ste20. | PMID: 12142009, PMID: 10970855 |
| YKL064W | MNR2 | NA | NA | NA | NA | -0.24 | 0 | Interacts with Fus3, which is downstream of Ste20. | PMID: 16319894 |
| YKR092C | SRP40 | NA | NA | NA | NA | -0.14 | 0 | Downregulated by pheromone in the Kar4 knockout mutant. | PMID: 17101777 |

| systematic name | standard name | eQTL FC | ptQTL FC | pQTL FC | phResQTL FC | phQTL FC | local QTL | Ste20 link | references |
|---|---|---|---|---|---|---|---|---|---|
| YLR006C | SSK1 | NA | NA | NA | NA | -0.07 | 0 | Co-regulator of the osmotic shock pathway, together with Ste20. | PMID: 10970855, PMID: 8808622 |
| YLR248W | RCK2 | NA | NA | NA | -0.54 | -0.15 | 0 | Direct target of Hog1. Interacts with Hog1, Cdc42, Msn2, Pbs2, and Ste50. | PMID: 10805732 |
| YLR335W | NUP2 | NA | NA | NA | NA | 0.28 | 0 | Direct phosphorylation target of Hog1, which is downstream kinase of ste20, and directly involved in gene expression regulation. | PMID: 23645671 |
| YLR399C | BDF1 | NA | NA | NA | NA | -0.17 | 0 | Bdf1 associates with the basal transcription factor TFIID, which has been linked to Ste20 and other genes of the Ste20 pathway. Bdf2 (paralog of Bdf1 and redundant) is phosphorylated by STE20. | PMID: 16118188, PMID: 16319894 |
| YML127W | RSC9 | NA | NA | NA | NA | -0.22 | 0 | Direct interactor of Hog1, which is a downstream kinase of Ste20 and directly involved in gene expression regulation. | PMID: 19153600 |
| YNL074C | MLF3 | NA | NA | NA | 0.71 | 0.21 | 0 | Interacts with Gic1/Gic2, which is downstream of Cdc42, a direct interactor of Ste20. | PMID: 16816427 |
| YOL070C | NBA1 | NA | NA | NA | NA | 0.13 | 0 | Interacts with Bem1, Cdc24, Boi1/2, and inhibits Cdc42, which are all components of the Ste20 pathway. | PMID: 25416945, PMID: 19841731, PMID: 28751498 |
| YOR014W | RTS1 | NA | NA | NA | NA | 0.29 | 0 | Involved in stress response in relation with Hog1, although there is no direct interaction with Hog1. Hog1 is downstream of Ste20. | PMID: 8846889 |
| YLR219W | MSC3 | NA | NA | NA | -0.39, -0.55 | NA | 0 | Differentially phosphorylated upon pheromones, therefore might be involved in the Gpa1/Ste20 pathway. | PMID: 21329885, PMID: 15665377 |

**Supplementary Table S10** Targets of the Gpa1/Ste20 hotspot on ChrVIII:1 which could be only indirectly or weakly linked to Ste20 and/or Gpa1 through literature research and are therefore not shown in Figure 5. These genes represent putative novel components of the Gpa1/Ste20 pathways.

| gene | name | eQTL FC | ptQTL FC | pQTL FC | phResQTL FC | phQTL FC | Local QTL | Ste20 link | references |
|---|---|---|---|---|---|---|---|---|---|
| YBR057C | MUM2 | 0.05 | NA | NA | NA | NA | 0 | Response to alpha-factor, i.e. component of pheromone response, and known to interact with Ste50. | PMID: 14585977, PMID: 22875988 |
| YHL006C | SHU1 | 0.15 | NA | NA | NA | NA | 0 | Regulated by Met4 whose expression is dependent of Hog1, a key effector of the MAPK pathway. Adjacent to Ste20 on the chromosome. | PMID: 19234305, PMID: 22438580, PMID: 17604854 |
| YHL009C | YAP3 | 0.13 | NA | NA | NA | NA | 1 | A Yap3 knockout showed decreased resistance to hyperosmotic stress, and decreased fitness in alkaline pH. Ste20 is also involved in the osmotic stress response (via Hog1). Yap3 genetically interacts with Msn2 and Pbs2 (Hog1 pathway), and regulates the transcription of Ste3. | PMID: 21127252, PMID: 17417638 |
| YHL003C | LAG1 | 0.11 | NA | NA | NA | NA | 1 | Adjacent to Ste20 on the chromosome. | NA |
| YHL008C | YHL008C | 0.56 | NA | NA | NA | NA | 1 | Adjacent to Ste20 on the chromosome. | NA |
| YLR433C | CNA1 | 0.11 | NA | NA | NA | NA | 0 | MATa Cna1 Cna2 double mutants were more sensitive than wild-type cells or either single mutant to growth arrest induced by the mating pheromone a factor and failed to resume growth during continuous exposure to a factor. | PMID: 1651503 |
| YMR179W | SPT21 | -0.13 | NA | NA | NA | NA | 0 | Spt10 and Spt21 contribute to silencing at the HML-alpha locus, which is responsible for mating type regulation upon pheromone stimuli. | PMID: 21057056 |
| YAL017W | PSK1 | NA | NA | NA | 1.13 | 0.3 | 0 | Negative genetic interaction with Ste20. | PMID: 19269370 |
| YDR145W | TAF12 | NA | NA | NA | NA | -0.24 | 0 | Subunit of the TFIID and SAGA complexes. TFIID has been linked to Stes20 and other genes of the Ste20 pathway. Involved in regulating sexual differentiation in fission yeast. | PMID: 16118188, PMID:29079657 |
| YER033C | ZRG8 | NA | NA | NA | NA | 0.25 | 0 | Indirect link: Zrg8p has been shown to be involved in the RAM signaling network essential for cell polarity. Similarly, Ste20 regulates MAP kinase pathways controlling cell polarity. | PMID: 24062589, PMID: 15972461 |
| YNR006W | VPS27 | NA | NA | NA | NA | 0.34 | 0 | Vps27 is in negative genetic interaction with Hog1, and its knockout changes sensitivity to salt, i.e. osmotic stress. | PMID: 20093466, PMID: 22282571, PMID: 27708008 |
| YOR355W | GDS1 | NA | NA | NA | NA | -0.18, -0.26 | 0 | Negative genetic interaction with Ste50 and Bem1 | PMID: 27708008, PMID: 20093466 |
| YBL032W | HEK2 | NA | NA | NA | 0.48 | NA | 0 | Potentially interacts with Ste20. | PMID: 18805955 |
| YMR086W | SEG1 | NA | NA | NA | -0.56 | NA | 0 | A component of eiosomes. Eisosomes were connected to the Ste20 pathway | PMID: 26359496 |
| YPL004C | LSP1 | NA | NA | NA | 0.69 | NA | 0 | A component of eiosomes. Eisosomes were connected to the Ste20 pathway | PMID: 26359496 |
| YMR031C | EIS1 | NA | NA | NA | 0.77 | 0.25 | 0 | A component of eiosomes. Eisosomes were connected to the Ste20 pathway | PMID: 26359496 |

**Supplementary Table S11** GO-enrichments of transcripts that are targeted by chrVIII:1.

| ontology | GO.ID | Term | Annotated | Significant | Expected | Pvalue |
|---|---|---|---|---|---|---|
| BP | GO:0000755 | cytogamy | 10 | 4 | 0.12 | 4.2E-06 |
| BP | GO:0000742 | karyogamy involved in conjugation with c... | 13 | 4 | 0.16 | 0.000014 |
| BP | GO:0000750 | pheromone-dependent signal transduction ... | 28 | 5 | 0.34 | 0.000019 |
| BP | GO:0000753 | cell morphogenesis involved in conjugati... | 14 | 3 | 0.17 | 0.00059 |
| BP | GO:2000220 | regulation of pseudohyphal growth | 18 | 3 | 0.22 | 0.00128 |
| BP | GO:0022604 | regulation of cell morphogenesis | 29 | 3 | 0.36 | 0.00212 |
| BP | GO:0000754 | adaptation of signaling pathway by respo... | 24 | 3 | 0.3 | 0.003 |
| BP | GO:0046020 | negative regulation of transcription fro... | 11 | 2 | 0.14 | 0.00765 |
| BP | GO:1900429 | negative regulation of filamentous growt... | 11 | 2 | 0.14 | 0.00765 |
| BP | GO:0043547 | positive regulation of GTPase activity | 105 | 5 | 1.29 | 0.00909 |
| BP | GO:0120031 | plasma membrane bounded cell projection ... | 12 | 2 | 0.15 | 0.0091 |
| MF | GO:0016836 | hydro-lyase activity | 28 | 3 | 0.34 | 0.0047 |
| MF | GO:0005096 | GTPase activator activity | 56 | 4 | 0.69 | 0.0048 |
| CC | GO:0043332 | mating projection tip | 116 | 7 | 1.43 | 0.0005 |
| CC | GO:0000131 | incipient cellular bud site | 64 | 4 | 0.79 | 0.0077 |
| CC | GO:0098797 | plasma membrane protein complex | 12 | 2 | 0.15 | 0.0091 |

**Supplementary Table S12** Targets of the Gpa1/Ste20 hotspot on ChrVIII:1 with no known link to neither Ste20 nor Gpa1, which are therefore not shown in Figure 5. These genes represent putative novel components of the Gpa1/Ste20 pathways.

| gene | name | eQTL FC | ptQTL FC | pQTL FC | phResQTL FC | phQTL FC | Local QTL | Function |
|---|---|---|---|---|---|---|---|---|
| YBR225W | YBR225W | 0.10 | NA | NA | NA | NA | 0 | Putative protein of unknown function; non-essential gene identified in a screen for mutants affected in mannosylphophorylation of cell wall components. |
| YDL059C | RAD59 | -0.14 | NA | NA | NA | NA | 0 | Protein involved DNA double-strand break repair. |
| YGL194C-A | YGL194C-A | 0.23 | NA | NA | NA | NA | 0 | Putative protein of unknown function. |
| YGR122C-A | YGR122C-A | 0.19 | NA | NA | NA | NA | 0 | Dubious open reading frame; unlikely to encode a functional protein, based on available experimental and comparative sequence data; similar to YLR334C and YOL106W. |
| YDR478W | SNM1 | -0.07 | NA | NA | NA | NA | 0 | Ribonuclease MRP complex subunit; ribonuclease (RNase) MRP cleaves pre-rRNA and has a role in cell cycle-regulated degradation of daughter cell-specific mRNAs. |
| YHL016C | DUR3 | 0.93 | NA | NA | NA | NA | 0 | Plasma membrane transporter for both urea and polyamines; expression is highly sensitive to nitrogen catabolite repression and induced by allophanate, the last intermediate of the allantoin degradative pathway. |
| YHR003C | TCD1 | -0.07 | NA | NA | NA | NA | 1 | tRNA threonylcarbamoyladenosine dehydratase; required for the ct6A tRNA base modification. |
| YHR067W | HTD2 | 0.08 | NA | NA | NA | NA | 0 | Mitochondrial 3-hydroxyacyl-thioester dehydratase. |
| YHR118C | ORC6 | -0.04 | NA | NA | NA | NA | 0 | Subunit of the origin recognition complex (ORC); ORC directs DNA replication by binding to replication origins and is also involved in transcriptional silencing; phosphorylated by Cdc28. |
| YKL135C | APL2 | 0.06 | NA | NA | NA | NA | 0 | Beta-adaptin subunit of the clathrin-associated protein (AP-1) complex; binds clathrin; involved in clathrin-dependent Golgi protein sorting; protein abundance increases in response to DNA replication stress. |
| YKL183W | LOT5 | -0.09 | NA | NA | NA | NA | 0 | Protein of unknown function. |
| YLR417W | VPS36 | 0.11 | NA | NA | NA | NA | 0 | Component of the ESCRT-II complex; contains the GLUE (GRAM Like Ubiquitin binding in EAP45) domain which is involved in interactions with ESCRT-I and ubiquitin-dependent sorting of proteins into the endosome. |
| YMR226C | YMR226C | 0.10 | NA | NA | NA | NA | 0 | NADP(+)-dependent serine dehydrogenase and carbonyl reductase; acts on serine, L-allo-threonine, and other 3-hydroxy acids. |
| YNL290W | RFC3 | -0.08 | NA | NA | NA | NA | 0 | Subunit of heteropentameric replication factor C (RF-C); which is a DNA binding protein and ATPase that acts as a clamp loader of the proliferating cell nuclear antigen (PCNA) processivity factor for DNA polymerases delta and epsilon. |
| YNR028W | CPR8 | -0.09 | NA | NA | NA | NA | 0 | Peptidyl-prolyl cis-trans isomerase (cyclophilin); catalyzes the cis-trans isomerization of peptide bonds N-terminal to proline residues. |

| gene | name | eQTL FC | ptQTL FC | pQTL FC | phResQTL FC | phQTL FC | Local QTL | Function |
|---|---|---|---|---|---|---|---|---|
| YNR032W | PPG1 | 0.08 | NA | NA | NA | NA | 0 | Putative serine/threonine protein phosphatase; putative phosphatase of the type 2A-like phosphatase family, required for glycogen accumulation. |
| YOL106W | YOL106W | 0.08 | NA | NA | NA | NA | 0 | Dubious open reading frame; unlikely to encode a functional protein, based on available experimental and comparative sequence data. |
| YOR180C | DCI1 | 0.12 | NA | NA | NA | NA | 0 | Peroxisomal protein; identification as a delta(3,5)-delta(2,4)-dienoyl-CoA isomerase involved in fatty acid metabolism is disputed. |
| YDR241W | BUD26 | 0.03 | NA | NA | NA | NA | 0 | Dubious open reading frame; unlikely to encode a functional protein, based on available experimental and comparative sequence data; not conserved in closely related Saccharomyces species; 1% of ORF overlaps the verified gene SNU56; diploid mutant displays a weak budding pattern phenotype in a systematic assay. |
| YGL060W | YBP2 | 0.13 | NA | NA | NA | NA | 0 | Central kinetochore associated protein; mediates mitotic progression; interacts with several central kinetochore proteins and centromeric histone Cse4p; role in resistance to oxidative stress; similar to Slk19p; YBP2 has a paralog, YBP1, that arose from the whole genome duplication. |
| YHL017W | YHL017W | 0.05 | NA | NA | NA | NA | 0 | Putative protein of unknown function; green fluorescent protein (GFP)-fusion protein co-localizes with clathrin-coated vesicles; YHL017W has a paralog, PTM1, that arose from the whole genome duplication. |
| YHL018W | MCO14 | 0.07 | NA | NA | NA | NA | 0 | Putative 4a-hydroxytetrahydrobiopterin dehydratase; green fluorescent protein (GFP)-fusion protein localizes to mitochondria and is induced in response to the DNA-damaging agent MMS. |
| YHL020C | OPI1 | -0.08 | NA | NA | NA | NA | 0 | Transcriptional regulator of a variety of genes; phosphorylation by protein kinase A stimulates Opi1 function in negative regulation of phospholipid biosynthetic genes; involved in telomere maintenance; null exhibits disrupted mitochondrial metabolism and low cardiolipin content, strongly correlated with overproduction of inositol; binds to phosphatidic acid. |
| YHL023C | NPR3 | 0.07 | NA | NA | NA | NA | 0 | Subunit of the Iml1/SEACIT complex; SEACIT (Iml1-Npr2-Npr3) is a subcomplex of SEAC, a coatomer-related complex that associates dynamically with the vacuole; Npr3 may have a structural or regulatory role, supporting Iml1p function as a GAP for the Rag family GTPase Gtr1, and leading to inhibition of TORC1 signaling in response to amino acid deprivation; SEACIT is required for non-nitrogen-starvation-induced autophagy; null mutant has meiotic defects; human NPRL3 homolog. |
| YHL025W | SNF6 | 0.04 | NA | NA | NA | NA | 0 | Subunit of the SWI/SNF chromatin remodeling complex; involved in transcriptional regulation; functions interdependently in transcriptional activation with Snf2 and Snf5; relocates to the cytosol under hypoxic conditions. |
| YKL044W | MMO1 | -0.15 | NA | NA | NA | NA | 0 | Protein of unknown function; SWAT-GFP, seamless-GFP and mCherry fusion proteins localize to the mitochondria; mRNA |

| gene | name | eQTL FC | ptQTL FC | pQTL FC | phResQTL FC | phQTL FC | Local QTL | Function |
|---|---|---|---|---|---|---|---|---|
| | | | | | | | | identified as translated by ribosome profiling data; MMO1 is a non-essential gene. |
| YML062C | MFT1 | 0.04 | NA | NA | NA | NA | 0 | Subunit of the THO complex; THO is a nuclear complex comprised of Hpr1, Mft1, Rlr1, and Thp2, that is involved in transcription elongation and mitotic recombination; involved in telomere maintenance. |
| YOR348C | PUT4 | -0.02 | NA | NA | NA | NA | 0 | Proline permease; required for high-affinity transport of proline; also transports the toxic proline analog azetidine-2-carboxylate (AzC); PUT4 transcription is repressed in ammonia-grown cells. |
| YPR127W | YPR127W | -0.12 | NA | NA | NA | NA | 0 | Putative pyridoxine 4-dehydrogenase; differentially expressed during alcoholic fermentation; expression activated by transcription factor YRM1/YOR172W; green fluorescent protein (GFP)-fusion protein localizes to both the cytoplasm and the nucleus. |
| YGR260W | TNA1 | NA | NA | -0.13 | NA | NA | 0 | High affinity nicotinic acid plasma membrane permease. |
| YKL114C | APN1 | NA | NA | -0.11 | NA | NA | 0 | Major apurinic/apyrimidinic endonuclease; 3'-repair diesterase. |
| YIL108W | YIL108W | NA | NA | -0.06 | NA | NA | 0 | Putative metalloendopeptidase; forms cytoplasmic foci upon DNA replication stress. |
| YLR144C | ACF2 | NA | NA | -0.08 | NA | NA | 0 | Intracellular beta-1,3-endoglucanase; expression is induced during sporulation; may have a role in cortical actin cytoskeleton assembly; protein abundance increases in response to DNA replication stress. |
| YCL057W | PRD1 | NA | NA | NA | 0.75 | 0.47 | 0 | Zinc metalloendopeptidase; found in the cytoplasm and intermembrane space of mitochondria. |
| YDR181C | SAS4 | NA | NA | NA | NA | 0.16 | 0 | Subunit of the SAS complex (Sas2, Sas4, Sas5); acetylates free histones and nucleosomes and regulates transcriptional silencing. |
| YDR293C | SSD1 | NA | NA | NA | NA | 0.15 | 0 | Translational repressor with a role in polar growth and wall integrity; regulated by Cbk1 phosphorylation to effect bud-specific translational control and localization of specific mRNAs; interacts with TOR pathway components; contains a functional N-terminal nuclear localization sequence and nucleocytoplasmic shuttling appears to be critical to Ssd1 function. |
| YDR432W | NPL3 | NA | NA | NA | NA | -0.22 | 0 | RNA-binding protein; promotes elongation, regulates termination, and carries poly(A) mRNA from nucleus to cytoplasm. |
| YER164W | CHD1 | NA | NA | NA | NA | 0.3 | 0 | Chromatin remodeler that regulates various aspects of transcription; acts in in conjunction with Isw1b to regulate chromatin structure and maintain chromatin integrity during transcription elongation by RNAP II by preventing trans-histone exchange over coding regions; contains a chromo domain, a helicase domain and a DNA-binding domain; component of both the SAGA and SLIK complexes. |
| YGL114W | YGL114W | NA | NA | NA | NA | 0.48 | 0 | Putative protein of unknown function. |
| YGR277C | CAB4 | NA | NA | NA | NA | 0.33 | 0 | Subunit of the CoA Sythesizing protein complex. |

| gene | name | eQTL FC | ptQTL FC | pQTL FC | phResQTL FC | phQTL FC | Local QTL | Function |
|---|---|---|---|---|---|---|---|---|
| YHR098C | SFB3 | NA | NA | NA | NA | 0.18 | 0 | Component of the Sec23p-Sfb3p heterodimer of the COPII vesicle coat. |
| YIL106W | MOB1 | NA | NA | NA | NA | 0.17 | 0 | CDK inhibitor and nuclear anchor. |
| YJL092W | SRS2 | NA | NA | NA | NA | -0.16 | 0 | DNA helicase and DNA-dependent ATPase; involved in DNA repair and checkpoint recovery, needed for proper timing of commitment to meiotic recombination and transition from Meiosis I to II transition. |
| YJR127C | RSF2 | NA | NA | NA | NA | -0.22 | 0 | Zinc finger protetin; involved in transcriptional control of both nuclear and mitochondrial genes, many of which specify products required for glycerol-based growth, respiration, and other functions; RSF2 has a paralog, TDA9, that arose from the whole genome duplication; relocalizes from nucleus to cytoplasm upon DNA replication stress. |
| YLL018C | DPS1 | NA | NA | NA | 0.68 | 0.12 | 0 | Aspartyl-tRNA synthetase, primarily cytoplasmic; homodimeric enzyme that catalyzes the specific aspartylation of tRNA(Asp). |
| YNR053C | NOG2 | NA | NA | NA | NA | -0.21 | 0 | Putative GTPase; associates with pre-60S ribosomal subunits in the nucleolus and is required for their nuclear export and maturation. |

### Supplementary Data

**Supplementary Data S1** Batch-membership, strain, and respective files from RNA-seq and SWATH-MS are specified for each culture. Some cultures that were characterized on the proteomic level did not undergo transcriptomic characterization.

**File: metadata.tsv**

**Supplementary Data S2** Transcript levels that were corrected for gene length, batch effects, library size are given for each culture.

**File: expressionLevels.tsv**

**Supplementary Data S3** The inherited allele is specified for all strains at 25590 variants. The reference and alternative alleles, as well as the locus, are also included.

**File: genotypeFull.tsv**

**Supplementary Data S4** Normalized and batch corrected abundances of proteins for each culture. Protein levels were computed from peptide levels as specified in the methods.

**File: proteinLevels.tsv**

**Supplementary Data S5** Normalized and batch corrected abundances of phosphopeptides for each culture. Sequences of phosphopeptides and proteins of origin are specified.

**File: phosphoPeptideLevels.tsv**

**Supplementary Data S6** Residuals computed from batch corrected and normalized protein and transcript abundances as specified in the methods.

**File: proteinResiduals.tsv**

**Supplementary Data S7** Residuals computed from batch corrected and normalized phosphopeptide and protein abundances as specified in the methods.

**File: phosphoPeptideResiduals.tsv**

**Supplementary Data S8** eQTL results. The location, target, and significance for each eQTL at FDR<10% are included. The QTL are numbered and some QTL correspond to multiple rows, as they are comprised of distinct regions that are in high LD.

**File: eQTL.tsv**

**Supplementary Data S9** ptQTL results. The location, target, and significance for each ptQTL at FDR<10% are included. The QTL are numbered and some QTL correspond to multiple rows, as they are comprised of distinct regions that are in high LD.

**File: ptQTL.tsv**

**Supplementary Data S10** pQTL results. The location, target, and significance for each pQTL at FDR<10% are included. The QTL are numbered and some QTL correspond to multiple rows, as they are comprised of distinct regions that are in high LD.

**File: pQTL.tsv**

**Supplementary Data S11** phResQTL results. The location, target, and significance for each phResQTL at FDR<10% are included. The QTL are numbered and some QTL correspond to multiple rows, as they are comprised of distinct regions that are in high LD.

**File: phResQTL.tsv**

**Supplementary Data S12** phQTL results. The location, target, and significance for each phQTL at FDR<10% are included. The QTL are numbered and some QTL correspond to multiple rows, as they are comprised of distinct regions that are in high LD.

**File: phQTL.tsv**

**Supplementary Data S13** Reduced genotype based on genotypeFull.tsv. Neighboring variants in complete LD were collapsed to 3593 marker regions.

**File: genotypeMapping.tsv**

**Supplementary Data S14** Quantitative predictors representing population structure. These predictors were used while mapping to correct for the effects of population structure and to avoid false positive associations.

**File: populationStructureCorvariates.tsv**
